# SLAPSHOT reveals rapid dynamics of extracellularly exposed proteome in response to calcium-activated plasma membrane phospholipid scrambling

**DOI:** 10.1101/2023.03.26.534250

**Authors:** Sami T. Tuomivaara, Chin Fen Teo, Yuh Nung Jan, Lily Y. Jan, Arun P. Wiita

## Abstract

To facilitate our understanding of the often rapid and nuanced dynamics of extracellularly exposed proteomes during signaling events, it is important to devise robust workflows affording fast time resolution without biases and confounding factors. Here, we present **S**urface-exposed protein **La**beling using **P**eroxida**S**e, **H**_2_**O**_2_, and **T**yramide-derivative (SLAPSHOT), to label extracellularly exposed proteins in a rapid, sensitive, and specific manner, while preserving cellular integrity. This experimentally simple and flexible method utilizes recombinant soluble APEX2 peroxidase that is applied to cells, thus circumventing biological perturbations, tedious engineering of tools and cells, and labeling biases. APEX2 neither requires metal cations for activity nor contains disulfide bonds, conferring versatility for a wide spectrum of experimental setups. We applied SLAPSHOT followed by quantitative mass spectrometry-based proteomics analysis to examine the immediate and extensive cell surface expansion and ensuing restorative membrane shedding upon the activation of Scott syndrome-linked TMEM16F, a ubiquitously expressed calcium-dependent phospholipid scramblase and ion channel. Time-course data ranging from one to thirty minutes of calcium stimulation using wild-type and TMEM16F deficient cells revealed intricate co-regulation of known protein families, including those in the integrin and ICAM families. Crucially, we identified proteins that are known to reside in intracellular organelles, including ER, as occupants of the freshly deposited membrane, and mitovesicles as an abundant component and contributor to the extracellularly exposed proteome. Our study not only provides the first accounts of the immediate consequences of calcium signaling on the extracellularly exposed proteome, but also presents a blueprint for the application of SLAPSHOT as a general approach for monitoring extracellularly exposed protein dynamics.

**Highlights:** An enzyme-driven method to tag extracellularly exposed proteins in an unbiased manner with a superior combination of temporal resolution, spatial specificity, and sensitivity

A general approach applicable to primary and scarce cells without involving cellular engineering

Short time scale proteome dynamics of Jurkat cells with and without TMEM16F revealed by SLAPSHOT coupled with quantitative mass spectrometry provide insights into phospholipid scrambling-mediated plasma membrane remodeling

## Introduction

The dynamics of plasma membrane are integral to all aspects of cellular physiology, including signaling, metabolism, and developmental trajectories. The diversity in these behaviors is promoted by proteins tasked with molecular transport, signal reception and transduction, cell-cell and cell-matrix interactions, as well as structural support and cellular shaping. Cells can accommodate rapid and sometimes drastic shifts at the plasma membrane, including membrane folding and unfolding as well as endo- and exocytotic processes that require fission or fusion of membranes. Many of these processes are associated with the redistribution of proteins and other molecules between compartments in response to intra- and extracellular stimuli (Stone et al., 2017; Enkavi et al., 2019; Jacobson et al., 2019; Marinko et al., 2019; Mao et al., 2021). Efforts to study proteome-wide effects of plasma membrane remodeling processes such as membrane growth and expansion have been hampered by a general lack of suitable experimental approaches (McCusker and Kellogg, 2012).

Monitoring proteome changes during plasma membrane remodeling poses several challenges for discovery-oriented or proteome-wide analyses. Difficulties arise both from the generally low abundance of transmembrane proteins and from their recalcitrance to biochemical analysis (Vit and Petrak, 2017), as well as from the often rapid pace of the accompanying proteomic changes (Bootman and Bultynck, 2020; Grecco et al., 2011). These challenges highlight the importance of technology development to supplement the current armament of chemical and enzymatic enrichment methods, especially when applied to signaling-mediated plasma membrane remodeling events. Here we report a labeling method, inspired by classical biochemistry and recently blossomed proximity labeling approaches, that can rapidly tag polypeptides exposed to the extracellular environment (**Figure 1A)**. Specifically, we make use of the soluble purified recombinant APEX2 enzyme, originally engineered from soybean ascorbate peroxidase (Martell et al., 2017). The peroxidase activity of this enzyme generates short-lived phenoxyl radicals from phenolic substrates, and effectively labels extracellularly exposed proteins when applied to cellular preparations. Our approach resembles recently reported PECSL (Li et al., 2021) that employs a soluble form of commercially available horseradish peroxidase (HRP) isolated and purified from plant tissue, but is strategically different from other peroxidase-mediated cell surface labeling methods where the enzyme is either engineered for surface expression (Li et al., 2020), or anchored to the cell surface via additional binding motifs (Kirkemo et al., 2022). In addition to its fast kinetics, APEX2 neither depends on calcium binding, nor contains disulfide bonds, rendering it a versatile tool for interrogating processes in the extracellular milieu.

**Figure 1.**
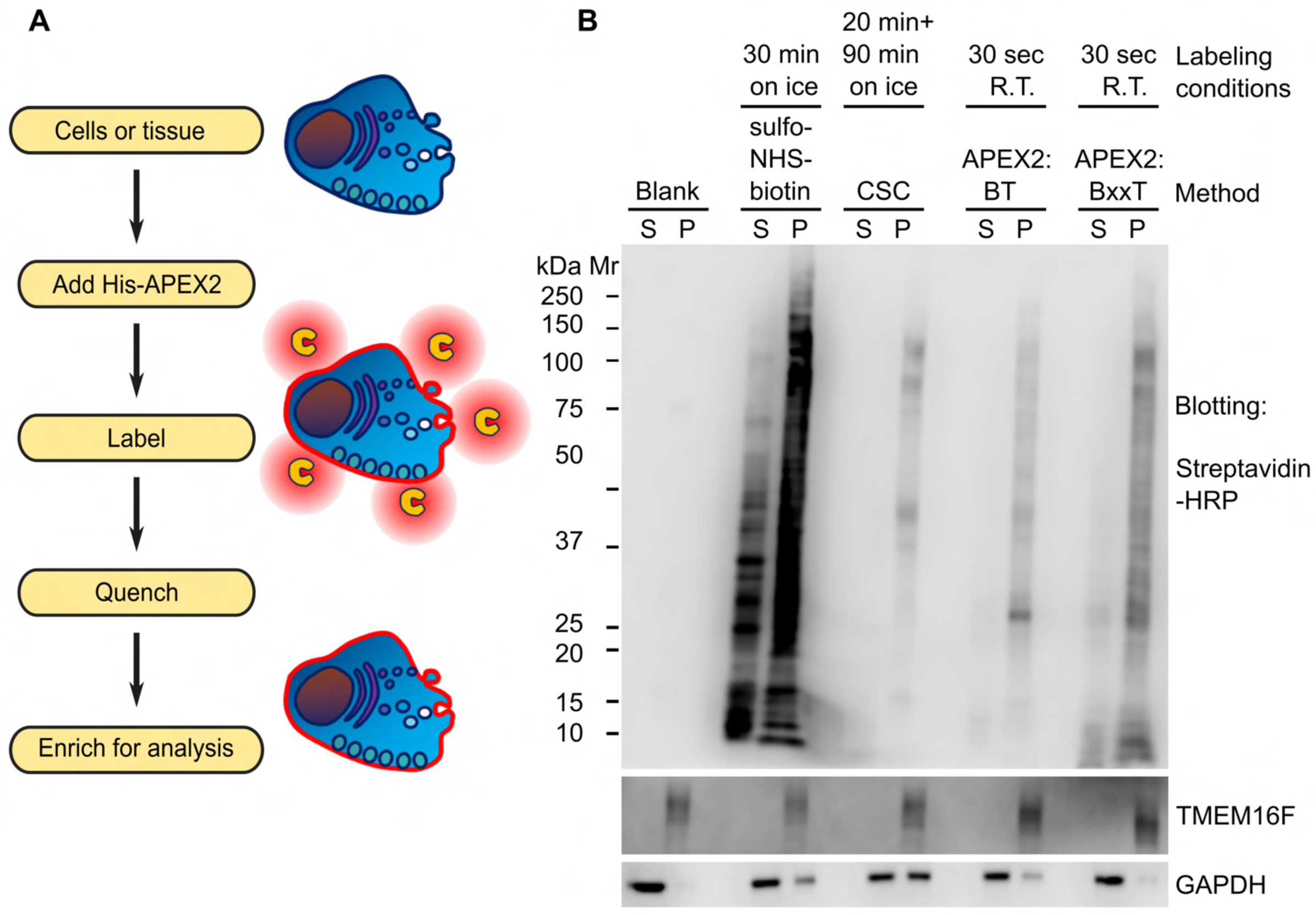
SLAPSHOT effectively labels extracellularly exposed proteins of cultured cells. (**A**) SLAPSHOT workflow. Purified soluble APEX2 enzyme is applied to the biological sample and the extracellularly exposed proteins are labeled in the presence of tyramide-derivative containing affinity handle such as biotin, and H_2_O_2_. Tagged proteins can be analyzed directly by streptavidin-blotting, or enriched for mass spectrometry or other analyses. (**B**) Streptavidin-blot analysis of U2OS cells labeled with sulfo-NHS-biotin, Cell Surface Capture (CSC), and SLAPSHOT. After labeling, the cells were hypotonically lysed, crudely fractionated into soluble and insoluble fractions by centrifugation, and the samples were probed with streptavidin-HRP. The blotting indicates spatial selectivity of CSC and SLAPSHOT, as nearly all of the biotin signal is segregated in the membrane fraction together with plasma membrane-residing TMEM16F protein, rather than with the soluble GAPDH. SLAPSHOT was performed using either cell permeable biotin-tyramide (BT) or cell-impermeable biotin-XX-tyramide (BxxT). Mr, relative molecular weight; kDa, kilodalton; S, soluble fraction; P, insoluble pellet fraction.

We rigorously optimized the experimental conditions, including reagent concentrations and labeling time, aiming for a robust and reproducible method that is widely applicable to address diverse biological questions. Toward this end, we also demonstrated that our method proceeds rapidly under chilled conditions, thus faithfully capturing the extracellularly exposed proteome dynamics associated with signaling events without interference from confounding factors such as endo- and exocytosis. Having established the powerful, accurate, and fast delivery of this method, we dubbed it SLAPSHOT for **S**urface exposed protein **La**beling using **P**eroxida**S**e, **H**_2_**O**_2_ and **T**yramide-derivative.

We applied SLAPSHOT to investigate a curious phenomenon where activation of the plasma membrane residing calcium-dependent phospholipid scramblase and non-selective ion channel TMEM16F elicits a rapid and massive Dynamin 2-mediated cell surface expansion (Bricogne et al., 2019; Deisl et al., 2021). Interestingly, cells lacking TMEM16F display a drastic reduction in surface area upon the same treatment (Bricogne et al., 2019). It is an intriguing open question regarding the origins of the recruited membrane for cell surface expansion and its proteomic composition, as well as the full physiological ramifications of this phenomenon. In addition to regulating cellular surface area, TMEM16F (Transmembrane Protein Family 16 Member F, encoded by the *ANO6* gene) has been shown to be a critical factor for phospholipid scrambling (Yang et al., 2012), extracellular vesicle release (Fujii et al., 2015), plasma membrane wound repair (Wu et al., 2020), syncytia formation (cell-cell fusion) for placenta formation during embryogenesis (Zhang et al., 2020) and as induced by SARS-CoV2 infection (Braga et al., 2021), as well as host cell entry of various pathogens including HIV (Zaitseva et al., 2017), Ebola virus (Acciani et al., 2021), and SARS-CoV2 (Sim et al., 2022). Relating to its role in extracellular vesicle release, TMEM16F is also the causative factor in Scott syndrome, a rare congenital bleeding disorder (Suzuki et al., 2010), where platelets of individuals with homozygous loss-of-function mutations fail to execute non-apoptotic plasma membrane phosphatidylserine scrambling necessary for the release of coagulating factor-containing extracellular vesicles (Ahmad et al., 1989; Bevers et al., 1992).

We monitored the extracellularly exposed proteomes of wild-type (WT) and TMEM16F CRISPR knock-out (16F KO) Jurkat cells in response to calcium stimulation using SLAPSHOT coupled to a quantitative mass spectrometry (MS) workflow. Our results revealed drastic differences in the extracellularly exposed proteomes of WT and 16F KO cells at their respective resting states, as well as differential responses to calcium stimulation. Bioinformatic analyses of SLAPSHOT-derived proteomics data provide hints on the plasma membrane and intracellular compartment dynamics during surface area expansion in WT cells as well as on the plasma membrane turnover in 16F KO cells during surface area reduction. Our data sheds light into this duality by revealing the dynamics of more than four thousand proteins, of which more than four hundred are membrane proteins, and their concerted actions in the presence and absence of TMEM16F activity. The data showcased here provide glimpses into the proteomic aspects of the plasma membrane architecture and dynamics, calcium signaling, and the interplay of cellular membranes. We report this study as an example of the usage of SLAPSHOT designed to empower investigations into the enigmatic and poorly understood nature of membrane biology.

## Results

### SLAPSHOT: A rapid enzymatic workflow for tagging extracellularly exposed proteins

To devise a method to rapidly tag global extracellularly exposed proteomes of living cells (**Figure 1A**), we subcloned, expressed, purified, and heme-reconstituted a soluble *N*-terminally His-tagged APEX2 protein (**Supplementary Figure S1**). We found that soluble APEX2 is stable in refrigerated conditions but loses activity when frozen (**Supplementary Figure S2A**). We systematically evaluated a series of labeling conditions to optimize the biotinylation levels on both adherent U2OS and suspended Jurkat cells, by using streptavidin-blotting (**Supplementary Figure S3**). In parallel, we developed a colorimetric microplate assay, based on the oxidation of catechol to ortho-quinone in the presence of H_2_O_2_, to gauge and standardize the peroxidase activity of APEX2 preparations (**Supplementary Figure S2B and S2C**). We tested the effects of several parameters, including the enzymatic activity of APEX2, the duration of the labeling, as well as the concentrations of biotin-xx-tyramide (BxxT) and H_2_O_2_, on the labeling intensity (**Supplementary Figure S3A-C**). We demonstrated that robust and satisfactory incorporation of biotin to adherent and suspended cells can be achieved in 30 and 45 seconds, respectively, at ambient temperature in the presence of 0.0005 AU s^−1^ µL^−1^ APEX2 activity (see **Materials and Methods**), 0.5 mM BxxT, and 0.5 mM H_2_O_2_. Throughout this work, we used the abovementioned specified enzymatic activity of APEX2, rather than its molar amount, for comparable labeling across biological samples and APEX2 preparations. Our results from the microplate-based optimization of H_2_O_2_ concentration (**Supplementary Figure S2D**) are consistent with the streptavidin-blotting assays (**Supplementary Figure S3B**) as well as with the extensive original characterizations of the APEX2 enzyme performed by the Ting lab (Lam et al., 2015). Using the colorimetric assay as well as a microplate-based turbidity assay (**Supplementary Figure S2A and E**), we also concluded that the introduction of water-soluble 10 mM Na-azide and 10 mM Na-ascorbate can fully quench the labeling reaction.

We then examined the spatial specificity of APEX2 labeling on U2OS cells using either biotin-tyramide (BT) or the membrane-impermeable biotin-xx-tyramide (BxxT) as substrates in parallel with two established surface biotinylation approaches: sulfo-NHS-biotin (Hurley et al., 1985) and the Cell Surface Capture (CSC) (Wollscheid et al., 2009). After hypotonically lysing and fractionating the denucleated cellular components into the particulate fraction of crude membranes and the cytosolic fraction, we probed the fractions with streptavidin-blotting. As shown in **Figure 1B**, biotinylated materials from APEX2 labeling were nearly completely segregated in the insoluble membrane-containing fraction that also contained the established plasma membrane protein TMEM16F (Suzuki et al., 2010; Yang et al., 2012) as revealed by western blot. These data indicate high spatial specificity of SLAPSHOT.

### SLAPSHOT aids in probing the extracellularly exposed proteomes without compromising cellular integrity

Next, we performed a proof-of-principle proteomics analysis on U2OS cells comparing SLAPSHOT to CSC (Wollscheid et al., 2009; Bausch-Fluck et al., 2015; van Oostrum et al., 2019), which has achieved the gold standard status for cell surfaceome analyses. Using label-free proteomics approach, we collected data from five replicate sets, each including cells treated with the two methods, as well as a non-treated control. To fairly compare the results, the same fraction of total peptides from each sample were analyzed. We subtracted the Intensity-Based Absolute Quantitation (iBAQ) values of the controls from those of the SLAPSHOT and CSC-labeled samples to obtain intensities free of contributions from the non-specifically bound materials to the solid support. The total tally of identified proteins stood at 3602 (**Figure 2A, Supplementary Table 1**) when proteins with iBAQ value greater than zero in at least one SLAPSHOT or one CSC sample were included in the analysis. Of these proteins, 1513 (42.0% of total) were detected by both methods, 183 (5.1%) and 1906 (52.9%) were detected exclusively by CSC and SLAPSHOT, respectively.

**Figure 2.**
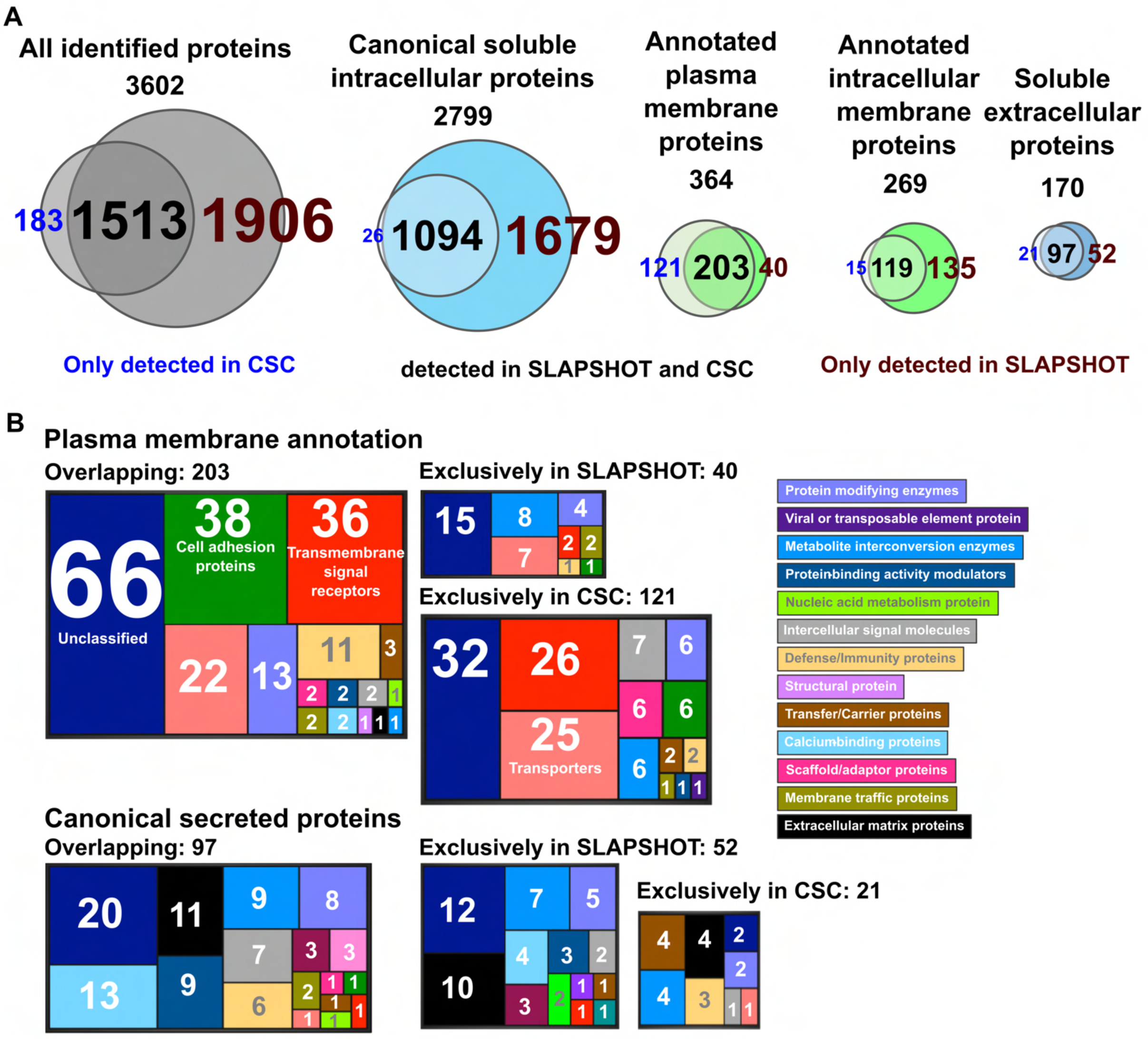
The numbers and annotations of identified proteins in U2OS cells using SLAPSHOT and CSC. (**A**) Venn diagrams indicate the number of identified proteins, in total and with various localization annotations. In each diagram, the proteins identified by CSC are on the *left* and proteins identified by SLAPSHOT are on the *right*. (**B**) PANTHER protein classifications. Total number of proteins, as well as proteins annotated in various compartments, are shown.

If one considers only proteins with plasma membrane (364 in total) and canonical secreted (170 in total) annotations, the percentage of overlap increases slightly to 55.8% (203 IDs) and 57.1% (97 IDs), respectively (**Figure 2A**). Of proteins detected exclusively by CSC, 121 are plasma membrane-annotated and 21 are secreted, whereas of proteins detected exclusively by SLAPSHOT, 40 are plasma membrane-annotated and 52 are secreted. Of the 1814 proteins without plasma membrane or secreted annotation that were exclusively detected by SLAPSHOT, 1679 are annotated as soluble intracellular proteins, and 135 are transmembrane proteins localized to various organelles. Notably, these 1679 proteins with intracellular annotation include 1509 (89.9%) that have been detected in the secretome and/or the extracellular vesicle proteome of U2OS cells (Jerez et al., 2017), or have mouse homologs present in the mitovesicles (extracellular vesicles that contain mostly mitochondrially derived proteins) (D’Acunzo et al., 2021). PANTHER protein class categorization of the identified plasma membrane and secreted proteins reveal functional classes expected from their localizations, including cell adhesion, transmembrane signal reception and transduction, as well as extracellular matrix scaffolding (**Figure 2B**). These data suggest that the analytical depth of SLAPSHOT in capturing extracellularly exposed proteins is complementary to that of CSC. Hierarchical clustering (**Supplementary Figure S4**) supports this notion as the samples cluster strongly based on the workflow used.

Given the abundant labeling of proteins annotated as intracellular by SLAPSHOT, we further investigated contributions from potential methodological artifacts using microscopy. First, we examined the integrity of the U2OS cell membrane after ambient temperature SLAPSHOT using a commercial Live-or-Dye reagent. As shown in **Supplementary Figure S5A**, the intracellular staining profile after SLAPSHOT was indistinguishable from that of the untreated negative control cells but drastically different from that of ethanol-perforated cells exhibiting strong intracellular staining, indicating that SLAPSHOT preserves the integrity of plasma membrane. Next, we directly examined the spatial selectivity of sulfo-NHS-biotin, CSC, and SLAPSHOT by labeling Jurkat cells, followed by visualization of the biotin distribution using fluorescently-conjugated streptavidin. The results presented in **Supplementary Figure S5B** indicate that biotinylation by all three methods was restricted to the cellular periphery, albeit with differing staining intensities. Altogether, these microscopic investigations revealed no contributions from methodological artifacts to the labeled proteome, in agreement with the biochemically obtained results. The low staining intensity observed in SLAPSHOT (**Supplementary Figure S5B**) compared to both sulfo-NHS-biotin and CSC results from the combination of short labeling time (45 s) as well as the low availability of the phenoxyl radical targets in the proteome. As revealed by our MS data, SLAPSHOT labels only tyrosine and tryptophan residues (**Supplementary Figure S6**), whose collective abundance as well as the fraction exposed to solvent is lower than those of lysines that are targeted by NHS-chemistry (Fiser and Simon, 2000; Sherchand et al., 2016). Furthermore, the microscopically detected CSC labeling intensity is inflated by the numerous and polymeric glycans decorating a large fraction of plasma membrane proteins (Schjoldager et al., 2020) as well as by the abundant plasma membrane glycolipids (Hanafusaet al., 2020). Taken together, these results demonstrate the feasibility and fidelity of SLAPSHOT labeling on living cells in shorter timescales than other technologies, without compromising the integrity of plasma membrane.

### Temporally resolved analyses of extracellularly exposed proteins from chilled cells to minimize vesicular traffic

Previous studies from the Hilgemann lab demonstrated that a brief 20 second calcium stimulation of wild-type (WT) Jurkat cells results in a rapid and massive cell surface area expansion, whereas TMEM16F-deficient Jurkat cells display an immediate reduction in the total cell surface area by “massive endocytosis” (MEND) as demonstrated by internalization of the membrane dye FM4-64 (Bricogne et al., 2019). The likely mechanistic control of the newly exposed membrane during cell expansion by Dynamin 2-dependent membrane unfolding was revealed in a recent follow-up study (Deisl et al., 2021). Little is known, however, about the proteomic composition of this newly deposited membrane. We surmised that SLAPSHOT could shed light on the TMEM16F-dependent membrane and extracellularly exposed proteome dynamics in this system. Before embarking on the analysis of the proteomic consequences of rapid membrane remodeling, we tested whether SLAPSHOT could also proceed in chilled conditions, under which vesicular trafficking and endo- and exocytotic processes are largely arrested (Tomoda et al., 1989; Walther et al., 2006). We established that labeling efficiencies comparable to those at the ambient temperature can be achieved by doubling the labeling times for adherent and suspended cells subjected to ice-cold conditions to 60 s and 90 s, respectively (**Supplementary Figure S7A**).

We proceeded to treat wild-type (WT) and TMEM16F knockout (16F KO) cells, generated using CRISPR-Cas9 technology (**Supplementary Figure S8A and 8B**), with 1 µM ionomycin for 1, 5, 10, or 30 min at 37 °C to facilitate extracellular calcium entry into the cytoplasm **(Supplementary Figure S9)**. At each stimulation endpoint, as well as for non-stimulated control (0 min, pre-stimulation) cells, an ice-chilled solution containing APEX2 and BxxT was added, followed by the addition of ice-chilled H_2_O_2_ to initiate the labeling. The reaction was allowed to proceed on ice for 90 s and quenched to ensure consistent labeling across samples. For each labeled sample, a corresponding negative control without BxxT was included. Biotinylated materials from the lysates of three replicate sets were enriched on a solid support, digested on-beads, and the resulting peptides were subjected to isobaric 10plex Tandem Mass Tag (TMT)-labeling (Werner et al., 2014), followed by quantitative LC-MS/MS analyses.

Collectively, we detected a total of 4148 proteins, as shown in volcano plots (**Figure 3A**, **Supplementary Figure S10**). At their resting states (pre-stimulation), we detected a total of 4138 proteins, of which 465 were exclusively detected in WT cells and 306 were exclusively detected in 16F KO cells. Of the 3360 proteins detected in both cell types at the resting state, only six displayed significant differences in extracellular exposure level between the cell types (absolute log_2_ fold change > 1, and *p*-value < 0.05, **Figure 3A**). Reactome analysis of the extracellularly exposed proteins in WT cells at resting state as well as 30 min post-ionomycin stimulation revealed an enrichment of proteins participating in vesicular trafficking, antigen presentation, and immune-signaling pathways (**Figure 3B)**. In contrast, extracellularly exposed proteins that were solely detected in the 16F KO cells are enriched in several predominantly intracellular processes, such as those related to mitochondrial homeostasis. Only a relatively small number of proteins displayed differential expression (*p*-value < 0.05) in response to calcium stimulation before the 30 min time point, compared to the non-stimulated control, in either cell type (**Supplementary Figure S10 and Supplementary Table 2**). Reproducibility of the multiplexed isobaric analyses is manifested by hierarchical clustering of the samples, whereby the individual samples segregate most strongly by the triplicate analyses performed, but not by timepoint or the cellular genotype (**Supplementary Figure S11**).

**Figure 3.**
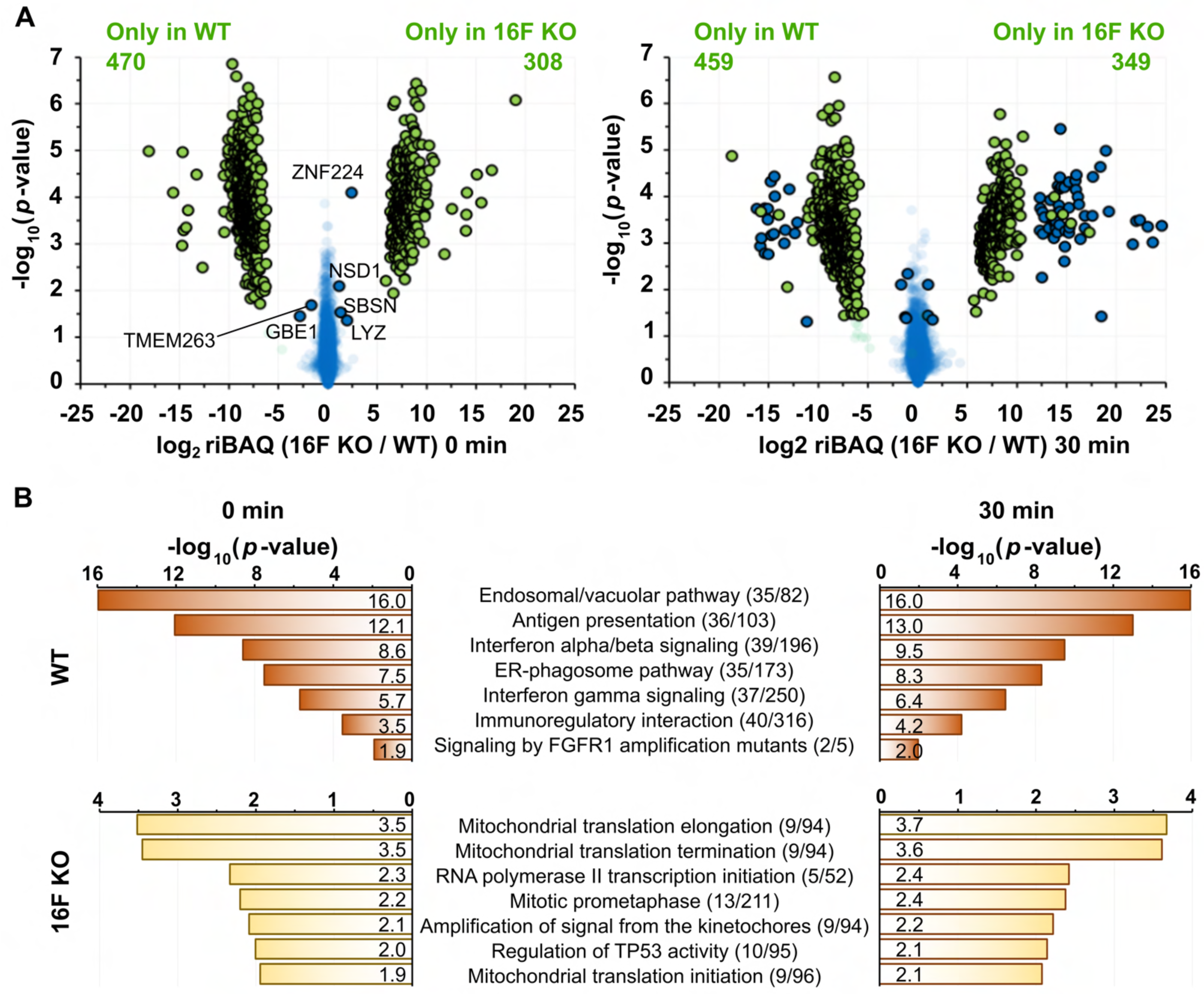
SLAPSHOT-identified proteins from WT and 16F KO Jurkat cells at resting state (*left panel*) and after 30 min of ionomycin stimulation (*right panel*). (**A**) Volcano plots depict the proteomics differences between these two cell types, for each cell type, *N* = 3. Blue datapoints indicate proteins that were identified in both samples. Datapoints in green were detected in only one of the samples and the other value was imputed. Circled datapoints have log_2_ transformed fold-difference of less than -1 or greater than 1 and are statistically significant (*p*-value < 0.05). (**B**) Reactome analysis of the proteins exclusively detected in the WT and 16F KO cells at the resting state and after 30 min of ionomycin stimulation. The pathways identified at 0 and 30 min are very similar in both cell types.

### TMEM16F-dependent dynamics of the extracellularly exposed proteome

To systematically investigate the trends and diversity in the time course profiles of the detected proteins, we fitted their riBAQ intensities against a collection of theoretical models (**Supplementary Table 3**) and classified each protein according to the best-fitting model. To simplify the model description, we assigned a single-letter code to represent the direction of the intensity change between two consecutive time points, such that “c”, “u”, and “d” respectively represent “constant”, “up”, and “down” (**Figure 4A**). For instance, a protein whose intensity decreases over the first interval (from 0 to 1 min) and increases over the other three intervals (1 to 5, 5 to 10, and 10 to 30 min) is classified as “d-u-u-u” (**Figure 4A**). Of the 61 theoretically computed patterns, 56 are represented by at least one detected protein (**Supplementary Figure S12**). Surprisingly, only 1167 (28.2%) out of the 4148 detected proteins occupy the same pattern in both cell types. Given that the normalized riBAQ values indicate the relative molar distribution of proteins both within a sample and between samples and hence could be used to appraise protein abundances, we tabulated the total riBAQ values for every time point in each pattern and visualized their overall distributions in a heatmap (**Supplementary Figure S12**).

**Figure 4.**
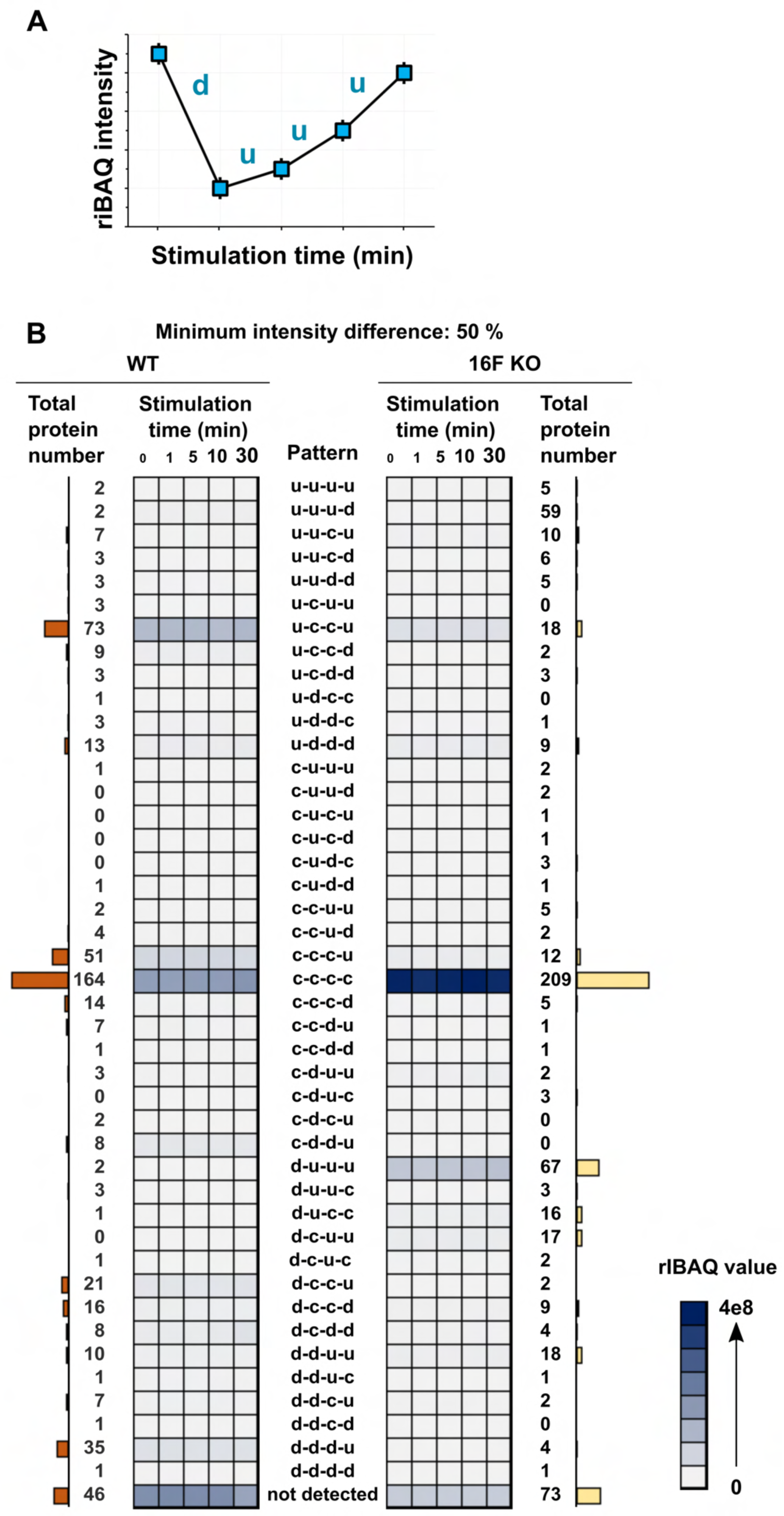
Proteins exhibit a wide variety of patterns in their extracellularly exposed levels in response to ionomycin stimulation. (**A**) An example of time-course intensity pattern for the extracellularly exposed protein with “d-u-u-u” expression profile where its intensity decreases between the first interval (from 0 to 1 min) and increases between the other three intervals (1 to 5, 5 to 10, and 10 to 30 min). (**B**) Heatmaps display the evolution of the total riBAQ for intensity patterns in WT (*left heatmap*) and 16F KO (*right heatmap*) cells, when considering proteins with membrane annotation (*N* = 533). Each square in the grid represents the sum of riBAQ values from the proteins with that intensity pattern. The total number of proteins belonging to a pattern from a given cell type are shown by bar graphs. Notably, most of the proteins as well as riBAQ intensity is concentrated in a few patterns.

As our impetus for developing SLAPSHOT was to identify and quantify the protein composition found on the rapidly expanding plasma membrane in response to TMEM16F activation, we performed analyses focusing solely on proteins with transmembrane and/or GPI-anchor domains (“membrane proteins”). Overall, a total of 488 and 460 membrane proteins were respectively identified in WT and 16F KO cells, populating 43 distinct patterns. Six patterns are prominent with at least 5% of all detected membrane protein IDs or 5% of the total riBAQ intensity (**Figure 4B**). We identified patterns where the fractional riBAQ intensity of the pattern (of total riBAQ intensity in the experiment) is either significantly smaller or significantly larger compared to the fraction of the identified membrane proteins in them (**Figure 5A**). A single pattern in WT cells, u-c-c-u, belongs to the former category: its 73 occupants, comprising 15% of the identified membrane proteins, contribute only 8.3% of the total riBAQ intensity, indicating that their molar amounts are approximately half that of the average detected membrane protein. Four patterns, c-c-c-u and d-d-d-u in WT cells, and d-u-u-u and c-c-u-u in 16F KO cells, exhibit the inverse trend, *i.e*., the membrane proteins in those patterns are present, on average, in higher molar amounts than the average membrane protein: 51 (10.5%) of c-c-c-u WT membrane proteins with 33.9% riBAQ, 35 (7.2%) of d-d-d-u WT membrane proteins with 14.9% riBAQ, 67 (14.6%) of d-u-u-u 16F KO membrane proteins with 34.5% riBAQ, and 5 (1.1%) c-c-u-u 16F KO membrane proteins with 7.8% riBAQ intensity.

**Figure 5.**
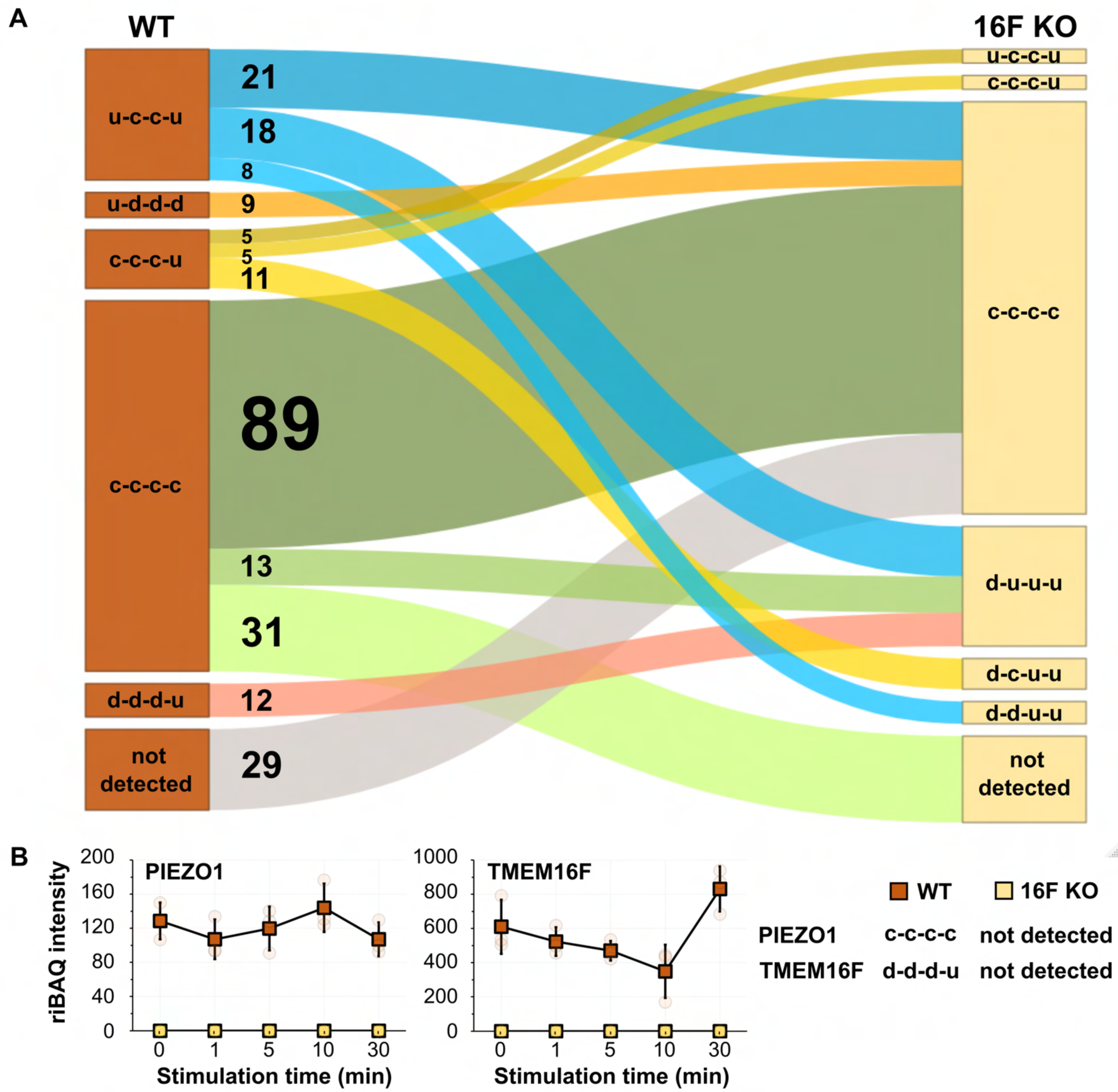
(**A**) Sankey flow diagram illustrates the shift of groups of membrane proteins from one intensity pattern in WT to another in 16F KO cells. Only the most prominent groups with number of proteins ≥ 5 (numbers indicated), and *p*-value < 0.05 are shown. WT u-c-c-u to 16F KO c-c-c-c, and WT c-c-c-c to 16F KO d-u-u-u contain less proteins than expected (based on hypergeometric distribution, *p*-value < 0.05), whereas all other transitions contain more proteins than expected. (**B**) Intensity time-course patterns of PIEZO1 and TMEM16F in response to calcium stimulation.

Diving into the protein constituents of these patterns, we gained the following insights:

a. Proteins with constant intensity (c-c-c-c pattern) are the largest group of membrane proteins in both cell types, with 164 (34%) and 209 (45%) proteins in WT and 16F KO cells, respectively, accounting for 31% and 39% of overall riBAQ intensities. Surprisingly, only 89 of the membrane proteins with this pattern are the same in both cell types, indicating that the membrane proteome dynamics undergo tremendous basal state alteration in the absence of TMEM16F.For instance, we found that the ubiquitous mechanosensitive calcium channel PIEZO1 is one of the 31 WT membrane proteins with c-c-c-c pattern that are not detected in 16F KO cells. Previous study has shown that raising intracellular calcium level via activation of PIEZO1 can activate TMEM16F which in turn leads to phospholipid scrambling and plasma membrane expansion to an extent similar to ionomycin stimulation (Bricogne et al., 2019). It is noteworthy that the extracellularly exposed PIEZO1 protein level in WT cells is very low as per its riBAQ value (**Figure 5B**). Due to the lack of commercially available anti-PIEZO1 antibodies that work unambiguously in Jurkat cell lysates, we examined the expression of PIEZO1 at the transcript level and found no difference between WT and 16F KO cells (**Supplementary Figure S13**), indicating that TMEM16F impinges on PIEZO1 surface expression at the protein level.
b. Membrane proteins from WT cells with d-d-d-u pattern include a group of canonical transmembrane plasma membrane proteins, such as those in the integrin and the intercellular adhesion molecular (ICAM) families (**Figure 6A**). Several members of the integrin and ICAM protein families are known to physically interact with each other (Staunton et al., 1988). Whereas they display d-d-d-u pattern on the WT plasma membrane during ionomycin stimulation, they are also co-regulated in the 16F KO cells but with the d-u-u-u pattern (see below), further suggesting a tight regulatory dynamic of this cluster of proteins. As canonical extracellular vesicle budding from the plasma membrane is TMEM16F activation-dependent (Ehlen et al., 2013; Fujii et al., 2015; Headland et al., 2015; Han et al., 2019; Wu et al., 2020), we speculate that membrane proteins with the d-d-d-u pattern in WT cells are budded from the plasma membrane into extracellular vesicles over the first ten minutes of the ionomycin stimulation (Bricogne et al., 2019) but the minuscule size of these microvesicles prevents them from sedimentation during the low-speed centrifugation employed to harvest the cells post-labeling. The rise at 30 min time-point of the membrane proteins with d-d-d-u pattern reflects their restoration along with general plasma membrane homeostasis from intracellular reserves. This projection is bolstered by the fact that TMEM16F itself is one of the proteins found in this pattern for WT cells. As expected, we detected TMEM16F in WT cells but not in the 16F KO cells (**Figure 5B**).
c. Massive endocytosis (MEND), whereby parts of the plasma membrane are acutely internalized upon calcium stimulation, has been previously demonstrated to be the primary membrane remodeling process of 16F KO Jurkat cells that do not release microvesicles (Bricogne et al., 2019). We speculate that 16F KO membrane proteins with d-u-u-u pattern (**Figure 6B**) are the major proteomic contributors to this phenomenon for several reasons. First, the immediate reduction in the protein intensity, followed by their restoration, matches the time-course and physical description of MEND. Second, d-u-u-u is the second largest membrane protein pattern in the 16F KO cells (after c-c-c-c), based on both the number of IDs and total riBAQ intensity. Their size contribution is also consistent with MEND where a large portion of plasma membrane is internalized, rendering proteins, including those in the integrin and ICAM families, inaccessible for SLAPSHOT. Furthermore, just two proteins with d-u-u-u pattern were detected in the WT cells while neither displayed the d-u-u-u pattern in 16F KO cells, also consistent with the uniqueness of MEND to the 16F KO cells. Finally, PANTHER analysis (GO cellular component) also supports this hypothesis, as the major localization annotations of these proteins are plasma membrane-associated (**Supplementary Figure S14**).
d. In reminiscence of the observation that treating WT Jurkat cells with calcium results in a rapid and massive cell surface area expansion (Bricogne et al., 2019), we speculate that the majority of the proteins that are freshly deposited along with the newly added membrane do so in a coordinated manner and can be accounted by the u-c-c-u pattern, for the following reasons. First, for WT cells, u-c-c-u is the only pattern with an immediate increase in intensity that contains a significant number of constituent membrane proteins (**Figure 4B**). Second, PANTHER analysis (using GO cellular component annotations) of these proteins indicates an annotated residency in intracellular membranes, including ER (**Supplementary Figure S14**). Third, their collective riBAQ value is significantly lower than the average membrane protein (8.3% of total riBAQ versus 15.0% of protein IDs), indicating their plasma membrane residency is not constitutive but triggered by ionomycin stimulation. These data are also consistent with Bricogne and co-workers’ observations whereby the newly deposited membrane is nearly devoid of electrophysiological activity and likely contains much less protein than the resting plasma membrane (Bricogne et al., 2019). Fourth, for the majority of the proteins that display the u-c-c-u pattern in the WT cells, their levels are either reduced or detectable but displaying the c-c-c-c pattern in 16F KO cells that do not display membrane expansion (**Figure 5C**).
e. The c-c-c-u pattern in the WT cells is the second most populated by the number of proteins, and the most populated by the total riBAQ intensity (**Figure 4B**). This group contains roughly equal numbers of ER and plasma membrane proteins, and curiously, mitochondrial proteins as well (**Figure 6D**). The increase of ER and plasma membrane proteins could be explained by the membrane reshuffling event while the cells are reverting to their resting state after TMEM16F-mediated phospholipid scrambling that causes extracellular vesicle shedding as well as the transient Dynamin 2-mediated membrane unfolding. The majority of the mitochondrial proteins detected in our assay are also found in mitovesicles, a novel class of extracellular vesicles (D’Acunzo et al., 2021). Mitovesicles consist mostly of mitochondrially-derived proteins and are surrounded by a double membrane akin to mitochondria themselves. Our SLAPSHOT labeling data suggests that mitovesicle release does not follow the trajectory of canonical extracellular vesicles whose release depends on TMEM16F activation, as indicated by the following observations (see below).
f. Curiously the c-c-u-u pattern in 16F KO cells only contains five proteins but collectively accounts for higher riBAQ value than many other patterns with more protein constituents (**Figure 6E**). The major contributor to the riBAQ value is VDAC3 (voltage-dependent anion-selective channel protein 3), also a constituent of mitovesicles (D’Acunzo et al., 2021). Prompted by this finding, we stratified the overall riBAQ values of all time points based on cellular compartment and found that proteins from mitovesicles are heavily present throughout the experimental duration (**Supplementary Figure S15**). Furthermore, we observed that in WT cells, their riBAQ intensities are relatively constant following ionomycin stimulation until a striking increase at the 30 min time point. In 16F KO cells, on the other hand, total riBAQ level of mitovesicle proteins increased slightly at a much earlier time point. Such phenomenon further confirms that mitovesicle release is insensitive to acute intracellular calcium elevation and does not depend on TMEM16F activity. However, the lack of TMEM16F on the plasma membrane may compromise the molecular machinery that regulate the release of mitovesicles.

**Figure 6.**
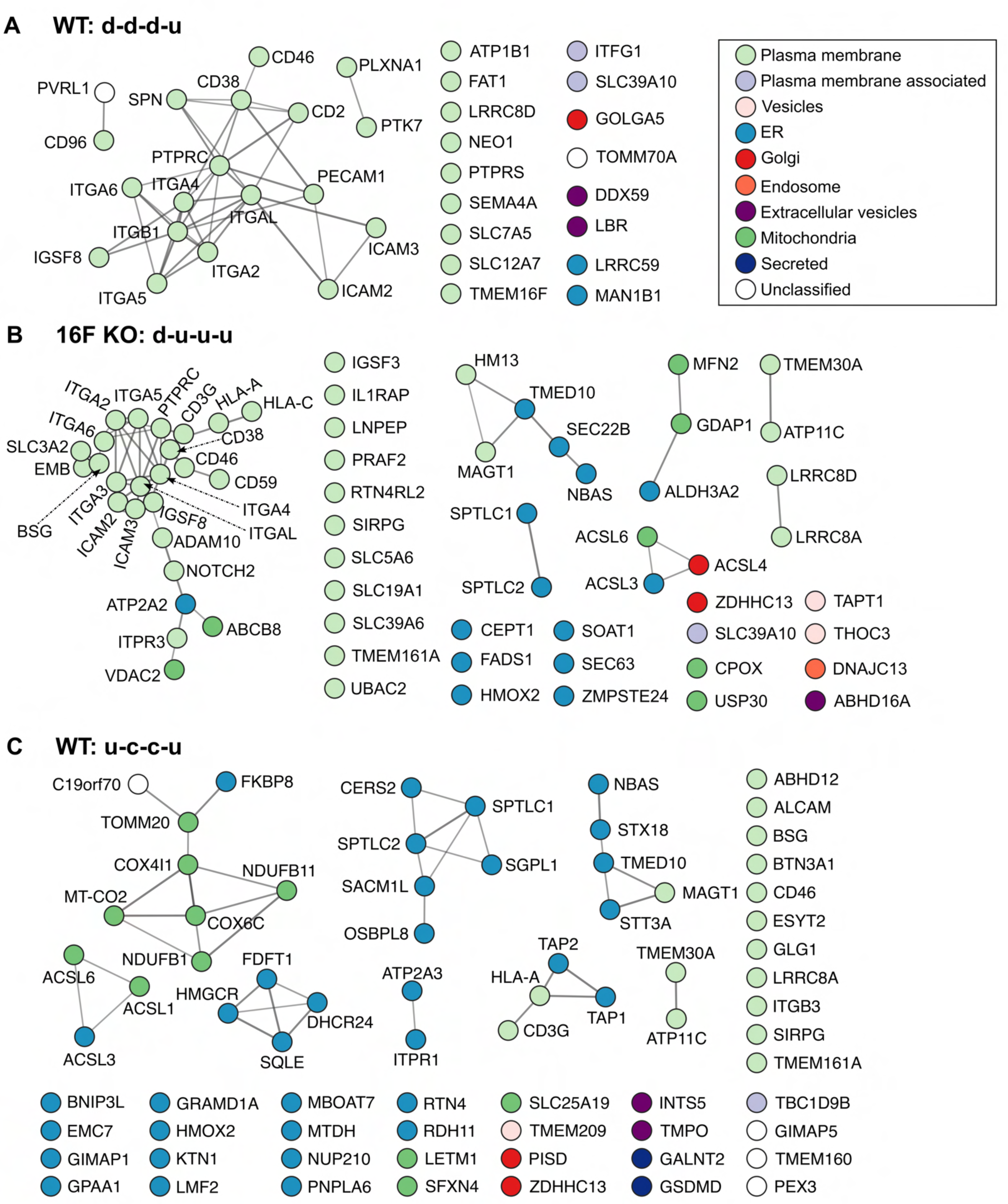
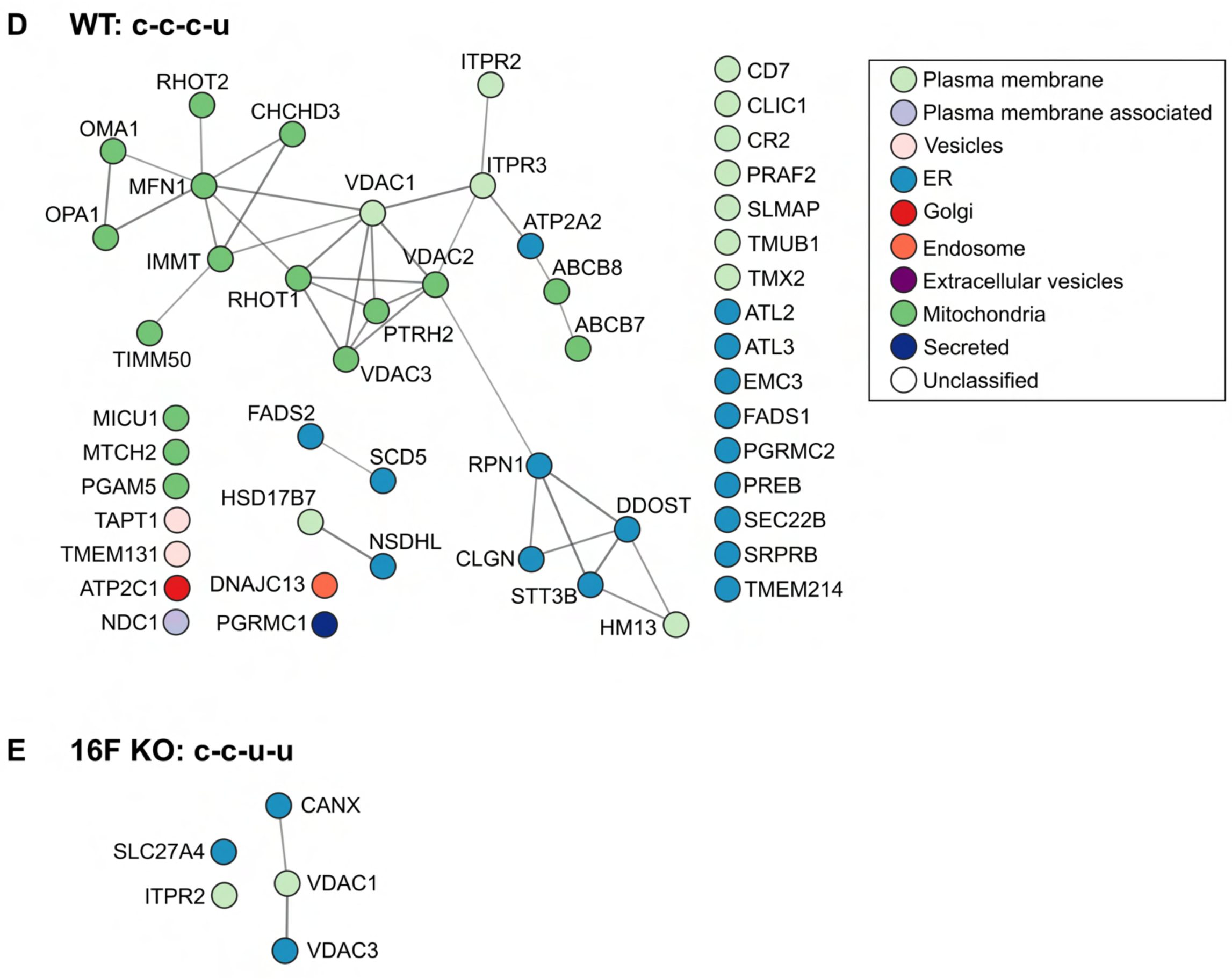
STRING analysis of proteins sharing a time-course intensity pattern. (**A**) WT d-d-d-u. (**B**) 16F KO d-u-u-u. (**C**) WT u-c-c-u. (**D**) WT c-c-c-u. (**E**) 16F KO c-c-u-u. While Plasma membrane proteins dominate most of these patterns, many ER and/or mitovesicle proteins also contribute to WT u-c-c-u and WT c-c-c-u patterns.

## Discussion

In this study we present SLAPSHOT, a protein tagging approach that utilizes purified soluble APEX2 enzyme to rapidly label extracellularly exposed proteins in cell culture models. After extensive optimizations, we conclude that SLAPSHOT compares favorably with the CSC technology (Bausch-Fluck et al., 2015) and sulfo-NHS-biotin labeling (Hurley et al., 1985), two widely used chemistry-based surface biotinylation methods. First, the extremely rapid peroxidase enzyme kinetics allows the capture of ephemeral processes typical for signaling at the plasma membrane. Protein labeling with such time-resolution is not feasible with the currently available chemistry-based methods, including CSC whose total workflow duration approaches two hours. Second, the sole APEX2 targets (Tyr and Trp sidechains), albeit at relative low abundance on proteins, are preferrable to labeling Lys residues or glycan post-translational modifications that could compromise workflow efficiency. For example, biotinylation of Lys residues by sulfo-NHS-biotin impedes digestion by proteases acting at this residue (Pauwels et al., 2021). In the case of CSC technology, ubiquitously found glycolipids on the plasma membrane actively compete for the reagent intended for glycoproteins, and for the enrichment capacity of the solid support (Wilchek and Bayer, 1987). Third, direct sidechain-tagging by the peroxidase enzymes accurately reflects the changes in the protein abundances, without relying on unpredictable proxy measurements whose fluctuations between samples may mask or completely overwhelm any underlying proteome changes. Aberrant glycosylation, for instance, often accompanies cellular pathological conditions (Frenkel-Pinter et al., 2017; Varki et al., 2015). Other post-translational modification, such as protein carbonylation resulting from oxidative stress, is also a direct target for hydrazide moiety (Akagawa, 2021), although its abundance on plasma membrane proteins has not been established. Escape altogether from the labeling by CSC is also a possibility for proteins exposed to the extracellular milieu given that some proteins, regardless of the secretory route, may remain non-glycosylated.

SLAPSHOT workflow also presents improvements compared to other hitherto published peroxidase-based surfaceome labeling methods. Unlike the recently reported approaches where APEX2 is physically tethered to the plasma membrane through an engineered binder (Kirkemo et al., 2022) or where HRP is fused to a plasma membrane anchored scaffold (Li et al., 2020), the soluble APEX2 employed in SLAPSHOT is free of any spatial constraints. Hence, SLAPSHOT does not introduce any protein or membrane microdomain-specific biases, and it can also effectively capture proteins normally residing intracellularly that are actively being exposed to the extracellular milieu regardless of their release mechanisms (**Figure 2, Supplementary Figure S5B**) while preserving cellular integrity (**Supplementary Figure S5A**). It is noteworthy that we store our APEX2 preparations refrigerated since we found significant loss of its activity when frozen (**Supplementary Figure S2A**). These findings can explain the suboptimal performance of this enzyme reported previously (Kirkemo et al., 2022).

SLAPSHOT does not require tedious and repetitive manipulations for adaptation to various biological models, and thus can be used for rare and primary cells or possibly *ex vivo* tissue samples that are innately refractive to genetic manipulations. Additionally, unlike the HRP utilized in PECSL (Li et al., 2021), APEX2 is neither metal-dependent nor redox-sensitive. Importantly, we established that the labeling efficiency of SLAPSHOT is not prohibitively hampered by ice-cold conditions (**Supplementary Figure S7A**), rendering it ideal for investigating proteome dynamics without confounding effects from cellular processes such as endocytosis. We have not performed cell number titrations to assay the minimum number of cells required by SLAPSHOT-MS, but data collected either from U2OS cells from one 10 cm dish, or from 5 million suspended Jurkat cells, with peptide material to spare, demonstrates its sensitivity and potential applications in labeling low abundance cell populations.

We applied SLAPSHOT to interrogate TMEM16F-dependent plasma membrane remodeling upon calcium stimulation in WT and TMEM16F KO Jurkat cells. We demonstrate that coupling SLAPSHOT with a quantitative proteomics workflow enables reconstruction of the extracellularly exposed status of more than four thousand proteins, of which more than four hundred are membrane proteins, after calcium stimulations as short as 1 min. We detected APEX2 protein in all Jurkat cell samples by MS. It is noteworthy that APEX2 riBAQ intensities are on average merely 2.5-times higher in the labeled samples compared to the corresponding non-labeled controls (**Supplementary Figure S16**). We attribute this observation to the visible colloid that forms during all APEX2 reactions, and that may contain polymerized BT/BxxT cross-linked with APEX2. We further note that the APEX2 intensities do not differ significantly among the non-stimulated and any of the calcium stimulated samples (**Supplementary Figure S16**). These data further lend credence to the notion that the membrane-impermeable reagents, short labeling time, and chilled conditions nearly completely preclude cellular uptake of APEX2 and other reagents by calcium-dependent endocytosis or any other processes. Collectively, our MS and other data indicate that SLAPSHOT complements existing labeling approaches and in many ways is applicable to wider experimental contexts.

In order to lay foundations on greater understanding of the effects of calcium signaling to plasma membrane dynamics, we aimed to identify and quantify proteins that are deposited to, or removed from, plasma membrane in response to the calcium-dependent activation of TMEM16F. Previous studies using Jurkat cells demonstrated that a combination of ionomycin and extracellular calcium elicits an instantaneous expansion of the cellular surface area in WT cells and that the same stimulation leads to a reduction of the cellular surface area in 16F KO cells (Bricogne et al., 2019; Deisl et al., 2021). By applying SLAPSHOT-coupled quantitative proteomic workflow on these cells after calcium stimulation, we systematically monitored the intensity profiles of 533 membrane proteins among 4148 extracellularly exposed proteins to reconstruct the plasma membrane landscape. At the resting state without calcium stimulation, we detected hundreds of proteins that are exclusively found on WT but not 16F KO cells, with several of those involved in antigen presentation and T cell receptor signaling (**Figure 3**). The absence of these proteins in 16F KO cells supports the notion that TMEM16F influences T cell receptor-mediated signaling as previously shown by others (Hu et al., 2016; Connolly et al., 2021). Moreover, the time course and riBAQ classification of all proteins detected in both WT and 16F KO cells revealed that the extracellularly exposed status of more than 86% of them have different intensity profiles between the cell types following calcium stimulation (**Supplementary Figure S11**). These data highlight the regulatory fluidity of the global protein network and the potentially complex effects upon the removal of the product(s) of just a single gene from the biological system, underscoring the need for a more systematic view of cellular dynamics.

Commensurate to their genotypes, we detected TMEM16F in the WT cells, but not in the 16F KO cells (**Figure 5B**). Due to the lack of flow cytometry- and microscopy-compatible anti-TMEM16F antibodies, there have been no facile approaches to examine its surface expression. By SLAPSHOT-MS, we captured the surface dynamics of TMEM16F in response to calcium stimulation, revealing a gradual decrease in surface levels before recovering to its pre-stimulation level (**Figure 5B**), a pattern that may reflect its intimate involvement with extracellular vesicle release (Ehlen et al., 2013; Fujii et al., 2015; Headland et al., 2015; Han et al., 2019; Wu et al., 2020). Others have shown that TMEM16F activation and subsequent cellular surface expansion are coupled with hypotonic medium-mediated calcium influx through PIEZO1 channel activity (Bricogne et al., 2019). We did not detect PIEZO1 in 16F KO cells by MS (**Figure 5A**), even though we did detect its transcript at comparable levels to that in WT cells (**Supplementary Figure S12**). Further studies are needed to decipher regulatory feedback mechanisms between TMEM16F and PIEZO1.

The role of calcium signaling in cellular physiology generally, and membrane physiology specifically, is well appreciated (Islam, 2020), but not much is known about its effects on the plasma membrane proteome dynamics. For example, in WT and 16F KO Jurkat cells overexpressing PD-1, Bricogne and co-workers used flow cytometry to detect the change in several membrane proteins, including transferrin receptor, CD28 and various HLA isoforms after 15 minutes of calcium stimulation. In our Jurkat cells that do not bear any transgene, we were able to detect several of these proteins by MS after SLAPSHOT labeling and reveal their dynamics following calcium stimulation (**Supplementary Figure S16**).

Focusing on the classification patterns of membrane proteins opens the avenue for deciphering protein contributions to the cellular events previously recorded via other methods, namely the TMEM16F-triggered plasma membrane expansion in WT cells and the massive endocytosis (MEND) observed in 16F KO cells. In WT cells, we identified a subset of proteins from the ER whose time course expression pattern (u-d-d-d) suggests that they correspond to the protein constituents residing on the unfolded membrane released by Dynamin 2 (**Figure 6A**, **Supplementary Figure S13A**). ER and plasma membrane contact sites are well known to participate in cellular calcium signaling and homeostasis (Crul and Maléth, 2021; Stefan, 2020). Here we provide new evidence that proteins with ER localization annotation can be exposed to the extracellular milieu upon calcium stimulation during the membrane unfolding process.

In 16F KO cells, we identified many plasma membrane proteins with d-u-u-u pattern suggesting their removal from the cell surface via endocytosis shortly following calcium stimulation (**Figure 6B**, **Supplementary Figure S13B**). Furthermore, we found that a cluster of cell adhesion proteins from the integrin and ICAM families exhibit identical d-d-d-u surface expression in WT cells following calcium stimulation but display the d-u-u-u pattern in 16F KO cells (**Figure 5A**, **Figure 6A** and **6B**). This scenario provides an example of the interconnectivity in the protein regulatory networks and showcases the effectiveness of the SLAPSHOT-MS approach in capturing biological information in an unbiased and systematic manner.

To our surprise, we detected a large number of mitochondrial proteins in our SLAPSHOT-MS data. Most of these proteins have been also found in mitovesicles, a novel type of extracellular vesicles whose release mechanism remains elusive but could be affected in pathological conditions (D’Acunzo et al., 2021). Results from our time course analysis suggest that the release of mitovesicles, unlike that of the canonical extracellular vesicles, does not depend on TMEM16F activation (**Supplementary Figure S15**). However, this pathway displays drastically different temporal patterns in WT and 16F KO cells, suggesting that altered membrane dynamics in these cell types may indirectly affect the mitovesicle release. Constitutive exocytosis (Stalder and Gershlick, 2020) and extracellular vesicle release (Aasebø et al., 2021) occurring without specific stimuli have also been demonstrated to be a fundamental cellular processes. Consistent with these scenarios, our initial MS analysis detected more proteins with intracellular annotation that presumably don’t contain glycosylations in SLAPSHOT-labeled compared to CSC-labeled U2OS cells, as well as a unique set of proteins with plasma membrane annotation that were not detected by CSC (**Figure 2**). We suggest that the large fraction of intracellularly annotated proteins in our MS data reflects the abovementioned or other poorly understood processes whose proteomic effects may have been underappreciated thus far due to technical limitations.

SLAPSHOT-MS yields quantitative and reproducible data that has high time-resolution, and that doesn’t require genetic engineering of the target cells, individualized “mono-plex” detection tools, or prior knowledge of the whereabouts of cellular proteins. Others have pondered dispatching MS data validation efforts by orthogonal detection methods such as western blotting (Aebersold et al., 2013; Mehta et al., 2022). In light of the technical maturity of quantitative MS-based proteomics, and the fact that immunoaffinity-based fluorescence-assisted cell sorting and microscopy approaches that are often used for validating, or in lieu of, surface proteomics, do not offer the same combination of time-resolution and “high-plexity” than SLAPSHOT, we echo these notions.

To summarize, we present an enzyme-based tagging method for extracellularly exposed proteins with fast kinetics both at ambient temperature and on ice, a method that can be easily applied to monitor rapid proteomics responses in a systematic and unbiased manner. Due to its simplicity and flexibility, SLAPSHOT can be readily adapted for studies that examine the dynamic changes of extracellularly exposed proteins in various biological contexts. Intriguing prospects for SLAPSHOT include the determination of cell surface processes during various other signaling events (Figard and Sokac, 2014; Hu et al., 2016; Enkavi et al., 2019) and membrane wound healing (Wu et al., 2020).

## Materials and Methods

All reagents were purchased from Sigma-Aldrich unless otherwise noted.

### Cloning, heterologous expression, and purification of hexahistidine-APEX2

We amplified the open reading frame encoding APEX2 from APEX2-NLS plasmid (Addgene plasmid #49386, a kind gift from Dr. Alice Ting) with Q5 polymerase (New England Biolabs) and subcloned the amplicon onto PCR linearized pRESTc vector (Invitrogen) with In-Fusion assembly reagent (Takara Bio). The resulting plasmid, pRESTc-His_6_-APEX2, was confirmed with Sanger sequencing and transformed into T7 Express *lysY* competent *E. coli* for recombinant protein expression. A single colony picked from a carbenicillin (carb)-containing LB plate was grown in LB/carb broth for overnight (37 °C, 250 rpm) and subcultured into Terrific Broth/carb (37 °C, 250 rpm) to an OD_600_ of 0.8 to 1.2. His-APEX2 protein expression was induced with IPTG at 0.5 mM and the bacteria were harvested 2 to 3 h post-induction. We resuspended the bacterial pellet in 150 mM NaCl, EDTA-free HALT protease inhibitor cocktail (Thermo Scientific), 1 mM phenylmethylsulfonyl fluoride (PMSF), 1 mg/mL lysozyme, 50 mM Tris-HCl (pH 7.5) lysis buffer and incubated it at room temperature for 30 min with occasional vortexing. We then added MgCl_2_ to a final concentration of 10 mM and DNAse I (GoldBio, >500 Kunitz U/mg) to a final concentration of 1 mg/mL and incubated the lysate for an additional 15 min at room temperature. From here on, APEX2 samples were handled either at +4 °C or on ice. After clarifying the lysate by centrifugation (12,000*g*, 15 min), we mixed the supernatant with pre-washed high-density cobalt-agarose resin (GoldBio) for batch binding. After 1 h incubation with gentle mixing, the resin was poured into an empty fritted column. We washed the resin extensively with 150 mM NaCl, 10 mM imidazole, 50 mM Tris-HCl (pH 7.5) buffer, eluted resin-bound APEX2 with 150 mM NaCl, 500 mM imidazole, 50 mM Tris-HCl (pH 7.5) buffer and concentrated the eluate with a pre-washed 10 kDa MWCO Amicon Ultra spin filter (Millipore). The concentrated eluate was buffer-exchanged into 20 mM K-phosphate (pH 7.0) using a pre-washed PD10 size-exclusion column (Bio-Rad) using gravity flow. We followed a previously reported procedure (Cheek et al., 1999) to prepare saturated hemin solution, and mixed it with the APEX2 sample for 3 h on ice (from hereon, the sample was kept protected from light to prevent photodamage). We next applied the hemin reconstituted APEX2 to the pre-washed DEAE-Sepharose resin slurry (Cytiva) and incubated for 30 min. The slurry was washed extensively with 20 mM K-phosphate (pH 7.0) buffer followed by elution with 100 mM K-phosphate (pH 7.0) buffer. Finally, we concentrated the APEX2 preparation with a pre-washed 10 kDa MWCO Amicon Ultra spin filter, sterile filtered it through a 0.2 μm filter, and stored it at 4 °C protected from light. Final protein concentration was measured with a BCA assay (Thermo Scientific) using BSA as a standard, and its densitometric purity was determined in Fiji (Schindelin et al., 2012). In contrast to others’ reports (Kirkemo et al., 2022; Lam et al., 2015), we did not freeze the purified protein, as this destroyed the enzymatic activity in our hands, and utilized the refrigerated APEX2 within four weeks of purification.

### APEX2 catechol colorimetric activity assay

We mixed 96 μL of 1 M glycine, 100 mM Tris-HCl (pH 7.5) buffer with 2 μL of appropriately diluted APEX2 and 2 μL of freshly made aq. 100 mM catechol solution onto a 96-well microplate. To this solution, we mixed 100 μL of 1 mM H_2_O_2_, 1 M glycine, 100 mM Tris-HCl (pH 7.5) buffer, and immediately measured absorbance at 480 nm using a Synergy 2 or Synergy H4 microplate reader (BioTek), at 2 s intervals. The activity of the APEX2 solution equals the slope of the absorbance-time curve (AU s^−1^ (2 μL)^−1^) for the early linear portion where Pearson correlation ≥ 0.998 (usually no more than 30 s when concentrated APEX2 solutions were used). The activity of the original APEX2 preparation (AU s^−1^ μL^−1^) was calculated by dividing the slope by the volume used in the activity assay (typically 2 μL), and by multiplying by any APEX2 dilution factor. We ascertained the linearity of the activity assay using a serial dilution of APEX2 solution. We also tested the effects of H_2_O_2_ concentration and quenchers (Na-azide, Na-ascorbate) doped in the H_2_O_2_ solution, as well as storage conditions, on the APEX2 activity. Blank experiments were conducted by omitting ether catechol or H_2_O_2_. The enzymatic activity of refrigerated APEX2 preparations were measured each day before the labeling experiments.

### APEX2 turbidity activity assay

We mixed 97 μL of 100 mM Tris-HCl (pH 7.5) buffer with 2 μL of APEX2 and 1 μL of 100 mM Biotin-tyramide (BT, APExBIO) in DMSO onto a 96-well microplate. To this solution, we then mixed 100 μL of 1 mM H_2_O_2_, 100 mM Tris-HCl (pH 7.5) buffer, and immediately measured the optical density at 400 nm using a Synergy 2 or Synergy H4 microplate reader (BioTek), typically at 2 s intervals for 1 min. We also tested the effects of quenchers (Na-azide, Na-ascorbate, Trolox) by doping them in the H_2_O_2_ solution. Blank experiments were conducted by omitting either BT or H_2_O_2_.

### Cell culture

We cultured all cells in humidified incubators at 37 °C with 5% CO_2_. Adherent U2OS cells (from Cell and Genome Engineering Core, UCSF) were grown to ∼90% confluency in DMEM media (Corning) supplemented with 10% FBS (Axenia BioLogix) on 3.5 cm (for optimizing conditions), or 10 cm (for proteomics) dishes. The Jurkat E6.1 T cells (a gift from Dr. Arthur Weiss, UCSF) were maintained in suspension at 0.2e6 to 1e6 cells/mL cell density either in regular RPMI1640 media supplemented with 10% FBS (Atlanta Biologicals) (for optimizing conditions) or in advanced RPMI1640 medium (Life Technologies) supplemented with 2% in-house heat-treated (56 °C for 30 min) FBS (Atlanta Biologicals) and GlutaMAX (for proteomics). AMO1 cells (a gift from Dr. Christoph Driesen, Kantonsspital St. Gallen, Switzerland) in suspension were cultured at 1e6 to 2e6 cells/mL cell density in RPMI 1640 medium supplemented with 1% penicillin/streptomycin (ThermoFisher Scientific) and 20% FBS (Atlanta Biologicals). We confirmed mycoplasma negativity for all cultures prior to experiments (DigitalTest v2.0, biotool.com). Replicate experiments were performed on cells from different passages.

### Generation of Jurkat TMEM16F knock-out cell line

We created a clonal TMEM16F (encoded by the *ANO6* gene) CRISPR-knockout (KO) Jurkat cell line using the Alt-R^TM^ CRISPR-Cas9 system (Integrated DNA Technologies, IDT) by following the manufacturer’s protocol. Briefly, we annealed two crRNAs both targeting human ANO6 exon 6 (crRNA1 sequence: tgtaaaagtacacgc and crRNA2 sequence: ctgaaaaaccggtcc, 3 μL of 200 µM each, IDT) independently with tracrRNA (3 μL of 200 µM, IDT) and mixed each of the RNA complexes with purified Cas9-NLS (12.2 μL of 41 μM, QB3 Berkeley, University of California) to obtain two tubes of RNPs each with a final volume of 20 μL. We prepared 1e6 Jurkat cells by washing them twice with PBS, and resuspended cell pellet with 45 μL Nucleofector^TM^ Solution V (Lonza) and subsequently mixed in 10 μL Supplement (Lonza), 5 μL of 96 μM Alt-R Cas9 Electroporation Enhancer (IDT), 20 μL of crRNA1-containing RNP and 20 μL of crRNA2-containing RNP. We transferred the cells to a cuvette supplied by the Nucleofector reagent, chilled it on ice for 5 min, and performed electroporation with a Nucleofector I device (Lonza) using a preset program (X-001). We let the cells recover in RPMI1640 media supplemented with 10% FBS for several days prior to single-cell clonal selection. We performed isogenic clone selection by allowing cells to under gradual expansion from a 96-well plate (seeded with a single cell per well on average) to larger vessels to obtain sufficient cell numbers for subsequent genotype confirmation. We used western blot detection of TMEM16F by an in-house generated rabbit anti-TMEM16F polyclonal antibody (#3016) (Yang et al., 2012) to screen for clones that do not display TMEM16F antibody cross-reactivity. Next, we isolated genomic DNA from the clones that passed the first round of selection and performed TA cloning, and submitted the miniprep plasmids for Sanger sequencing. Briefly, we added 25 mM NaOH, 0.2 mM EDTA to cell pellets and incubated the samples at 98 °C for an hour, and neutralized with 55 mM Tris-HCl (pH 5.5) to obtain genomic DNA (gDNA). To amplify the targeted genomic fragment for Sanger sequencing, we mixed gDNA with a primer set that recognizes the region outside the pair of guide RNA sequences (forward: AGT TGG TTT CTT GTT CTG CGT T; reverse: ACT CTT GCT CTG GCT TGA TGA T) together with 2x GoTaq master mix (Promega) and commenced the reaction according to the manufacturer’s protocol. After confirming the presence of amplicon in the PCR product, we purified the amplicons and performed TA cloning using a kit (Invitrogen). Plasmids from 8-10 colonies from a single TA cloning reaction were cultured for miniprep and submitted for Sanger sequencing to confirm their genotype prior to further experimentation.

### RT-PCR analysis

We harvested WT and 16F KO Jurkat cells by centrifugation and extracted RNA using a RNeasy mini kit from Qiagen. Next, we prepared cDNA using a ProtoScript First Strand kit (NEB) and used 1 μL of cDNA for 10 μL of PCR using 2x GoTaq master mix (Promega) and 250 nM of forward and reverse primer sets with conditions recommended by the manufacturer. Primer sets were as followed: PIEZO1 forward: GCT GTA CCA GTA CCT GCT GTG; PIEZO1 reverse: CAG CCA GAA CAG GTA TCG GAA G; Actin forward: GAC ATG GAA GCC ATC ACA GAC; Actin reverse: AGA CCG TTC AGC TGG ATA TTA C. PCR products were analyzed on 2% agarose gel and visualized with the aid of GelGreen nucleic acid gel stain (Biotium).

### Optimization of SLAPSHOT labeling for adherent and suspended cells

U2OS cells were washed thrice with PBS *in situ* without lifting. Jurkat cells were washed twice with PBS with centrifugation (300*g*, 3 min). We added 0.5 mL of solution containing 2× APEX2/BxxT (APExBIO) in PBS (U2OS in 3.5 cm dish, and 1e5 of Jurkat cells) to the cells with gentle swirling for 10 to 15 s. We initiated the labeling reaction by adding an equal volume of 2× H_2_O_2_ in PBS. After desired labeling time with constant and gentle mixing, the reaction was quenched by adding an equal volume of 2× ice-cold quencher solution in PBS. The cells were washed twice with the quenching solution before harvesting or fixing. The labeling reaction parameters were optimized for U2OS and Jurkat cells to the following values, in the following order (all cited concentrations 1×): 0.5 mM BxxT concentration and APEX2 activity (all combinations of 0.125, 0.25, or 0.5 mM BxxT and 0.000125, 0.00025, or 0.0005 AU s^−1^ μL^−1^ APEX2, with 1 mM H_2_O_2_, 1 min labeling time, room temperature); H_2_O_2_ concentration (0.0625, 0.125, 0.25, 0.5, or 1 mM H_2_O_2_, with 0.5 mM BxxT and 0.0005 AU s^−1^ μL^−1^ APEX2); labeling time and temperature (10, 20, 30, 45, or 60 s at room temperature, and 30, 60, 90, and 120 s on ice, with 0.5 mM H_2_O_2_).

### Other labeling methods and hypotonic crude fractionation

We performed Sulfo-NHS-biotin (Pierce Thermo Scientific) labeling of U2OS cells on ice for 30 min according to the manufacturer’s protocol and the CSC labeling of U2OS cells as previously described (Wollscheid et al., 2009). We then executed a hypotonic lysis protocol to obtain crude soluble and membranous fractions from SLAPSHOT, Sulfo-NHS-biotin, CSC, and non-labeled control cells, for streptavidin and immunoblotting according to a previously described protocol (Teo and Wells, 2014).

### Streptavidin and immunoblotting

We added RIPA lysis buffer (50 mM Tris (pH 7.4), 150 mM NaCl, 1% NP-40, 0.5% Na-deoxycholate, 0.1% Na-dodecyl sulfate, 10 mM (tris(2-carboxyethyl)phosphine) (TCEP), 1 mM EDTA, EDTA-free HALT protease inhibitor cocktail, 10 mM Na-azide, 10 mM Na-ascorbate) to control and labeled cells, left them on ice for 15 min, and centrifuged (16,000*g*, 15 min, 4 °C). We mixed an equal portion of the lysate supernatant with 4× LDS sample buffer (Invitrogen) supplemented with 50 mM TCEP and heated them at 60 °C for 15 min. We carried out protein gel electrophoresis with Mini-PROTEAN TGX Stain-Free gels (Bio-Rad) and applied UV light (5 min, ChemiDoc Touch imaging system, Bio-Rad) to activate the stain-free labeling. We then transferred the gels to Immobilon-P membranes (EMD Millipore) with the Trans-Blot transfer system (Bio-Rad) using 25 mM Tris, 192 mM glycine, and 20% (v/v) ethanol transfer buffer. We then quantified the stain-free labeled total protein on the transferred membranes using the ChemiDoc Touch imaging system. We detected the biotin incorporation using streptavidin-HRP conjugate (Jackson ImmunoResearch) as previously reported (Teo and Wells, 2014). For TMEM16F immunoblotting, we blocked the membrane with 5% non-fat milk dissolved in TBST, incubated with anti-TMEM16F antibody (1:2,000 dilution, (Yang et al., 2012)) for overnight at 4 °C, and donkey anti-rabbit antibody conjugated with HRP (1:24,000, Jackson ImmunoResearch). To detect GAPDH, we used an anti-GAPDH-HRP antibody (1:60,000, Proteintech group) for overnight at 4 °C. We washed the membrane extensively with deionized water followed by TBST after each antibody incubation. We used a C-DiGit scanner (LI-COR) for chemiluminescence detection after incubating with an appropriate ECL substrate (either Pierce ECL or SuperSignal West Pico Plus, Thermo Scientific).

### Visualization of biotin signal with microscopy

We performed the labeling, fixing, washing, and staining steps on Jurkat cells with repetitive centrifugations (800*g*, 5 min) to exchange the solution and wash the cells. We labeled the cells with SLAPSHOT, Sulfo-NHS-biotin, and CSC according to the abovementioned protocols. We washed away the excess reagents, fixed the cells with 2% PFA in PBS at room temperature for 10 min, quenched with 50 mM ammonium chloride solution in PBS at room temperature for 15 min, performed blocking with 5% BSA in PBS, incubated cells with streptavidin-conjugated with Alexa Fluor 488 (Invitrogen) at a concentration of 2 μg/ml for overnight at 4 °C. We performed two to three washes with PBS between each step and counter-stained with Hoechst 33342 (1 μg/mL in PBS, Invitrogen) in the final step of washing. We mounted the cells with Fluoromount-G (Southern Biotech) on 18 mm coverslips. We acquired the Alexa Fluor 488 and Hoechst 33342 signals on a Leica SP8 microscope with a 63x/NA1.40 objective with the default lasers for each fluorophore. Images were exported to Fiji software (Schindelin et al., 2012) for further compilation. While data is not shown, we found that immunostaining with SLAPSHOT labeling on adherent U2OS cells consistently leads to extremely high background in addition to the hallmark plasma membrane staining, presumably from staining of the cell- or media-derived proteins deposited on the coverslips.

### Live-or-Dye staining

U2OS cells were grown on 12 mm coverslips and labeled with SLAPSHOT according to the described protocol. After washing the cells with the quencher solution, we added freshly diluted Live-or-Dye 488/515 reagent (1:1000, Biotium) in PBS and incubated the cells on ice for 30 min. The cells were washed once with PBS and fixed with 2% PFA in PBS at room temperature for 10 min. We then washed the cells with PBS twice, counter-stained with 1 μg/mL Hoechst 33342 (Invitrogen) in PBS for 5 min and mounted the cells with Fluoromount-G (Southern Biotech). For negative control, we added Live-or-Dye 488/515 working solution to the cells without any manipulations. For positive control, we treated the cells with 10% ethanol in PBS for 10 min at 37 °C before staining. We acquired the Live-or-Dye 488/515 and Hoechst 33342 signals on a Leica SP8 microscope with a 40x/NA1.30 objective with the default laser settings for each fluorophore. Images were exported to Fiji software (Schindelin et al., 2012) for compilation.

### Mass spectrometry analysis of method comparison samples from U2OS cells

U2OS cells were seeded at 1e6 cells per 10 cm dish and grown for 48 h. The plates were washed thrice and labeled with CSC or SLAPSHOT in a final volume of 3 mL with constant gentle swirling, or left as non-labeled controls. After quenching and washing, we added RIPA buffer without TCEP to each plate and incubated it on ice for 15 min. After scraping and collecting the lysates, samples were cleared by centrifugation, and the supernatants were transferred to fresh tubes, snap-frozen, and stored at -80 °C until subsequent Neutravidin pull-down and shotgun proteomic experiments.

### SLAPSHOT labeling of ionomycin stimulated Jurkat cells

We washed Jurkat cells thrice in warm PBS and aliquoted 5e6 cells in 2 mL of PBS containing 1.8 mM CaCl_2_ into a 15 mL conical tube. We then added ionomycin (Cayman Chemical) to a final concentration of 1 μM and continued incubation at 37 °C for either 1, 5, 10, or 30 minutes, or left the cells untreated as controls. After the stimulation endpoint, we added 0.5 mL of ice-cold solution containing 0.005 AU s^−1^ µL^−1^, 5 mM APEX2, and 5 mM BxxT (all 10×) in PBS, and after a few seconds of gentle mixing, 2.5 mL of ice-cold 1 mM H_2_O_2_ in PBS to initiate the labeling. For each labeling reaction, we also performed mock labeling where the BxxT was omitted. After 90 s of labeling on ice with gentle swirling, we quenched the labeling reaction by adding 5 mL of ice-cold 20 mM Na-azide, and 20 mM Na-ascorbate in PBS. We then washed the cells twice in the ice-cold quenching buffer and once in PBS with centrifugation (500*g*, 3 min, 4 °C) and snap-froze the well-drained cell pellets. We stored the cell pellets at -80 °C until lysis.

### Capture and on-bead digestion of biotinylated proteins for mass spectrometry

We incubated cleared cell lysates prepared in RIPA buffer without TCEP (see **Streptavidin and immunoblotting**) with (slurry volume) 125 μL (Jurkat) or 50 μL (U2OS) of pre-washed high-capacity NeutrAvidin-agarose beads (ThermoFisher) on the rotisserie for 30 min at 4 °C (all sample handling from this point on was performed at 4 °C or on ice until otherwise noted) and removed the unbound material by centrifugation (1000*g*, 5 min, 4 °C). One percent aliquots of the lysates and the unbound fractions were saved for streptavidin blot analysis to ensure proper titer of beads and quantitative capture of biotinylated proteins. The agarose beads were washed five times with the lysis buffer without quenchers (additionally, protease inhibitors were omitted starting from wash two), five times with 1 M NaCl, 50 mM Tris-HCl (pH 7.4) buffer, and five times with 1.5 M urea, 250 mM ammonium bicarbonate (digestion buffer). Each wash was performed using 1 mL of buffer for 5 min on rotisserie followed by centrifugation. We then reduced and alkylated the samples with 250 μL of 10 mM Tris(2-carboxyethyl)phosphine (TCEP), and 40 mM iodoacetamide (IAA) in the digestion buffer for 30 min at room temperature protected from light. After removing the soluble reagents, we digested the samples on-bead in 250 μL of the digestion buffer containing 0.1% (w/v) RapiGest SF surfactant (Waters), 1 mM CaCl_2_, and 1 μg of trypsin/Lys-C (Promega) on a rotisserie for 18 h at ambient temperature. After digestion, we collected the soluble materials and washed the beads twice with 250 μL of 10 mM ammonium bicarbonate and pooled all unbound materials.

### Peptide clean-up with solid-phase extraction

We acidified the digested samples with concentrated trifluoroacetic acid (TFA) and cleaned up the peptides with C18 SOLA Solid-Phase Extraction (SPE) columns (ThermoFisher). Briefly, we re-loaded the flow-through from the first loading, washed the column first with 1 mL of 0.1% TFA and then with 0.1% formic acid (FA), and eluted bound materials with 0.1% FA in 60% acetonitrile. Loading and elution steps were performed unaided (by gravity) whereas the washes were aided by gently applying air pressure via a syringe. The eluted materials were dried in vacuum, redissolved in water, and quantified spectrophotometrically by A_280_ measurement on NanoDrop (ThermoFisher) using BSA peptide mixture as a standard.

### TMT10plex isobaric labeling of Jurkat cell peptides

We labeled 10 µg of tryptic peptides from each experiment with a TMT10plex Isobaric Labeling Reagent kit (Thermo Scientific) (Zecha et al., 2019). Unlabeled (control) and labeled samples from the five time points of a single replicate comprised a set that was labeled with the ten individual reagents. After quenching the reaction with hydroxylamine, the samples from the set were pooled, dried, redissolved, and fractionated with a Pierce High pH Reversed-Phase Peptide Fractionation Kit (Thermo Scientific) into eight fractions according to the manufacturer’s instructions. We measured the peptide concentration from each dried fraction by NanoDrop. The resulting peptide mixtures were analyzed by LC-MS/MS.

### Analysis of the amino acid targets of the phenoxyl radicals

We harvested AMO1 cells by centrifugation (500*g*, 3 min) and washed twice in PBS. The cell pellet was dissolved in 7.5 M urea 250 mM ammonium bicarbonate buffer (∼10e7 cells/mL) with sonication. After clarification (16,100*g*, 15 min, +4 °C) and measuring protein concentration by BCA assay, the proteins were reduced and alkylated with TCEP and IAM. After diluting the sample with 250 mM ammonium bicarbonate to a final urea concentration of 1.5 M, CaCl_2_ was added to a final concentration of 1 mM, trypsin/Lys-C (Promega) and 0.1% (w/v) RapiGest SF surfactant (Waters) and trypsin/Lys-C to 1:50 protease: protein ratio. The digestion was allowed to proceed for 18 h at ambient temperature. We cleaned the digested peptides by SPE and quantified them by NanoDrop. For the analysis of the labeling targets of APEX2, we prepared two vials, each with 5 µg of the cleaned-up peptides, 0.0005 AU s^−1^ µL^−1^ APEX2, and 1 mM BT in a total volume of 5 µL in PBS. APEX2 labeling was then initiated in one of the vials with 5 µL of 1 mM H_2_O_2_ in PBS, whereas the other was left as control and received just PBS. After 45 s of incubation, 10 µL of quenching buffer with 20 mM Na-azide, and 20 mM Na-ascorbate in PBS was added to both vials. We applied the reaction mixtures to a 3000 MWCO spin filter (NanoSep, Pall) to remove APEX2 and the formed colloid, and depleted the remaining biotinylated materials from the filtrate with 25 µL (slurry volume) of Pierce streptavidin magnetic beads (Thermo Scientific). Peptides in the supernatants were cleaned-up with C18 ZipTips (EMD Millipore) prior to LC-MS/MS analysis.

### Liquid chromatography-tandem mass spectrometry analyses

We performed Liquid Chromatography-Tandem Mass Spectrometry (LC-MS/MS) analyses on a Thermo Scientific Q Exactive Plus Hybrid Quadrupole-Orbitrap mass spectrometer interfaced to a Dionex UltiMate 3000 UHPLC system with a NanoSpray Flex source. The MS instrument was operated in positive mode with spray voltage and heating capillary temperature set to 2200 V and 250 °C, respectively. We injected the peptides in 5 μL of aqueous 0.1% FA (Solvent A) into an Acclaim PepMap RSLC C18 column (75 μm × 150 mm, 2 μm particle size, 100 Å pore size, Thermo Scientific) kept at 40 °C. Chromatographic separations were achieved by an increase in 0.08% FA in 80% acetonitrile (Solvent B) with the following gradient: 0 to 15 min: 3% B at 500 nL/min; 15 to 210 min: 3 to 50% B at 200 nL/min, followed by column washes at 500 nL/min.

Survey (MS^1^) scans for data-dependent acquisition (DDA) were recorded with 350 – 1500 m/z (label-free samples) or 375 – 1400 *m*/*z* (TMT-labeled samples) scan range, 70,000 resolution (at 200 *m*/*z*), 1.7 *m*/*z* (label-free) or 0.7 *m*/*z* (TMT) isolation window, 3e6 AGC target, 100 ms (label-free) or 50 ms (TMT) maximum IT, 20 s (label-free) or 30s (TMT) dynamic exclusion, exclusion of isotopes, as well as unassigned and +1 charge states, preferred peptide matches, and 27 (label-free) or 32 (TMT) NCE. A maximum of 12 (label-free) or 15 (TMT) HCD fragment (MS^2^) scans were acquired with fixed mass 100 *m*/*z*, 17,500 (label-free) or 35,000 (TMT) resolution (at 200 *m*/*z*), 5e4 (label-free) or 1e5 (TMT) AGC target, 2e3 minimum AGC target, and 180 ms (label-free) or 100 ms (TMT) maximum IT. We saved all spectra in the profile mode.

### Proteomics data analysis

We analyzed the RAW data files containing LC-MS/MS spectra with MaxQuant (Cox and Mann, 2008) (v1.6.17.0) against the human proteome FASTA file containing the reviewed canonical and isoform entries (obtained from UniProt on 2020-04-02) as well as the His-APEX2 sequence. We used the following search parameters: Trypsin/P digestion, 2 max missed cleavages, oxidation (M) and acetyl (protein N-term) as variable modifications, and carbamidomethyl (C) as fixed modification. All other settings were left with their default values, including a 1% false discovery rate (FDR) at both peptide and protein levels. We invoked a match between runs for the TMT and peptide depletion experiments, and Intensity Based Absolute Quantification (iBAQ) for the optimization and TMT and experiments. Reverse hits, as well as potential contaminants and protein groups only identified by site, were removed from consideration.

To obtain protein intensities that can be compared both within and among samples, we implemented the relative iBAQ-TMT (riBAQ-TMT) quantitation scheme (Shin et al., 2013) for the TMT-labeled samples, whereby the TMT channel intensities for a given protein were scaled so that the sum from all channels equals the iBAQ value (Schwanhäusser et al., 2011). We then subtracted the intensities of non-labeled control channels from those of the corresponding labeled channels. Proteins with resulting zero or negative intensity were considered not detected in the sample. In the statistical R computing environment (R Core Team, 2021) (v4.0.5 or newer), we applied miceRanger (Wilson, 2021) (v1.3.4) to impute missing values, and internal reference scaling (IRS) normalization with pseudo-reference (Plubell et al., 2017) to correct batch effects. The imputation was however ignored and protein intensities were left to zero if the protein was not detected in any of the six samples (three labeled and three controls) from a given sample.

### Bioinformatics analyses

Differential protein expression (fold-change and *p*-value) was assessed by paired two-tailed Welch’s tests in Perseus (Tyanova et al., 2016) (v1.6.15.0) without multiple test adjustments. We utilized protein subcellular localization annotations and other information from UniProt (The UniProt Consortium, 2021), Human Protein Atlas (v19.3, *proteinatlas.org*) (Thul et al., 2017), VesiclePedia (v4.1) (Kalra et al., 2012), and Surface Protein Atlas (Bausch-Fluck et al., 2015). Pathway and gene set analyses were performed in PANTHER (v16) (Mi et al., 2021; Mi and Thomas, 2009), and Reactome (v79) (Gillespie et al., 2022). PANTHER analyses were performed with Fisher’s exact test without multiple test adjustments. Redundant and overlapping categories were removed. STRING analysis (Szklarczyk et al., 2021) (v11.5) in the Cytoscape environment (Shannon et al., 2003) (v. 3.9.1) was used to derive physical associations between proteins. A diverse set of theoretical model profiles (*N* = 2182) were constructed (**Supplementary Table 2)** that vary both in the directions and magnitudes of intensity changes between the time points. The initial model set was filtered to contain only those models (*N* = 2109) with at least 50% difference between smallest and largest value, in addition to the “static” model. Protein intensity profile distances to model profiles were calculated in R with the philentropy package (Drost, 2018) (v. 0.6.0) using Bhattacharyya distance and the protein was assigned a profile with the model with the smallest distance. After distance calculations, model descriptions were simplified by collapsing models with the same general pattern, regardless of the intensity information. Enrichment factor (E) of the overlapping proteins belonging to particular profiles in the WT and 16F KO cells was calculated with E = (n × N) / (wt × ko), where n is the number of overlapping proteins, N is the total number of detected proteins (4148), wt is the number of proteins with a particular profile in the WT cells, and ko is the number of proteins with a particular profile in the 16F KO cells. The *p*-values for the overrepresentation and underrepresentation of the overlaps were calculated with Fisher’s exact test in base R. Hierarchical clustering was performed in R with the ComplexHeatmap package (Gu et al., 2016) (v1.10.2) with method and metric set to single and Pearson correlation, respectively.

### Reagent availability

The plasmid encoding the soluble hexahistidine-APEX2 is available at Addgene (TO BE ADDED). TMEM16F KO Jurkat cell line is available from the Jan lab upon request.

### Data availability

The mass spectrometry proteomics data have been deposited to the ProteomeXchange Consortium (http://proteomecentral.proteomexchange.org) via the PRIDE partner repository (Vizcaíno et al., 2013) with the dataset identifier TO BE ADDED.

### Code Availability

All R code used to generate data in this paper is presented in the **Supplementary Materials**.

## Author Contributions

S.T.T. and C.F.T. conceived the project, executed all experiments, analyzed and interpreted the data, and wrote the manuscript. Y.N.J., L.Y.J., and A.P.W. oversaw the project and provided the resources for the project. All authors discussed and commented on the results and participated in editing the manuscript.

## Acknowledgements

We thank Drs. Wen Lu and Arthur Weiss for providing Jurkat E6.1 cells as well as their insight into Jurkat cell biology, Dr. Christoph Driesen for providing AMO1 cells, Dr. Alice Ting for providing APEX2-NLS plasmid, Drs. Mark Burlingame and Kamal Mandal for assistance with mass spectrometry, Drs. Ya-Chu Chang, David Crottès, John Gilchrist, and Han-Hsuan Liu for helpful suggestions and critical readings of the manuscript, and Jan and Wiita lab members for their input. C.F.T. is a recipient of a UCSF PBBR post-doctoral grant that sponsored part of this work. This study is supported by R35 NS122110 to L.Y.J. and DP2 OD022552 to A.P.W. Lily Jan and Yuh Nung Jan are investigators of the Howard Hughes Medical Institute.

