## Supplementary Figures for "SLAPSHOT reveals rapid dynamics of extracellularly exposed proteome in response to calcium-activated plasma membrane phospholipid scrambling"


| Figure | Page |
| --- | --- |
| Supplementary Figure S1 | 2 |
| Supplementary Figure S2 | 3 |
| Supplementary Figure S3 | 5 |
| Supplementary Figure S4 | 6 |
| Supplementary Figure S5 | 7 |
| Supplementary Figure S6 | 9 |
| Supplementary Figure S7 | 11 |
| Supplementary Figure S8 | 12 |
| Supplementary Figure S9 | 13 |
| Supplementary Figure S10 | 14 |
| Supplementary Figure S11 | 16 |
| Supplementary Figure S12 | 17 |
| Supplementary Figure S13 | 19 |
| Supplementary Figure S14 | 20 |
| Supplementary Figure S15 | 21 |
| Supplementary Figure S16 | 22 |
| Supplementary Figure S17 | 23 |

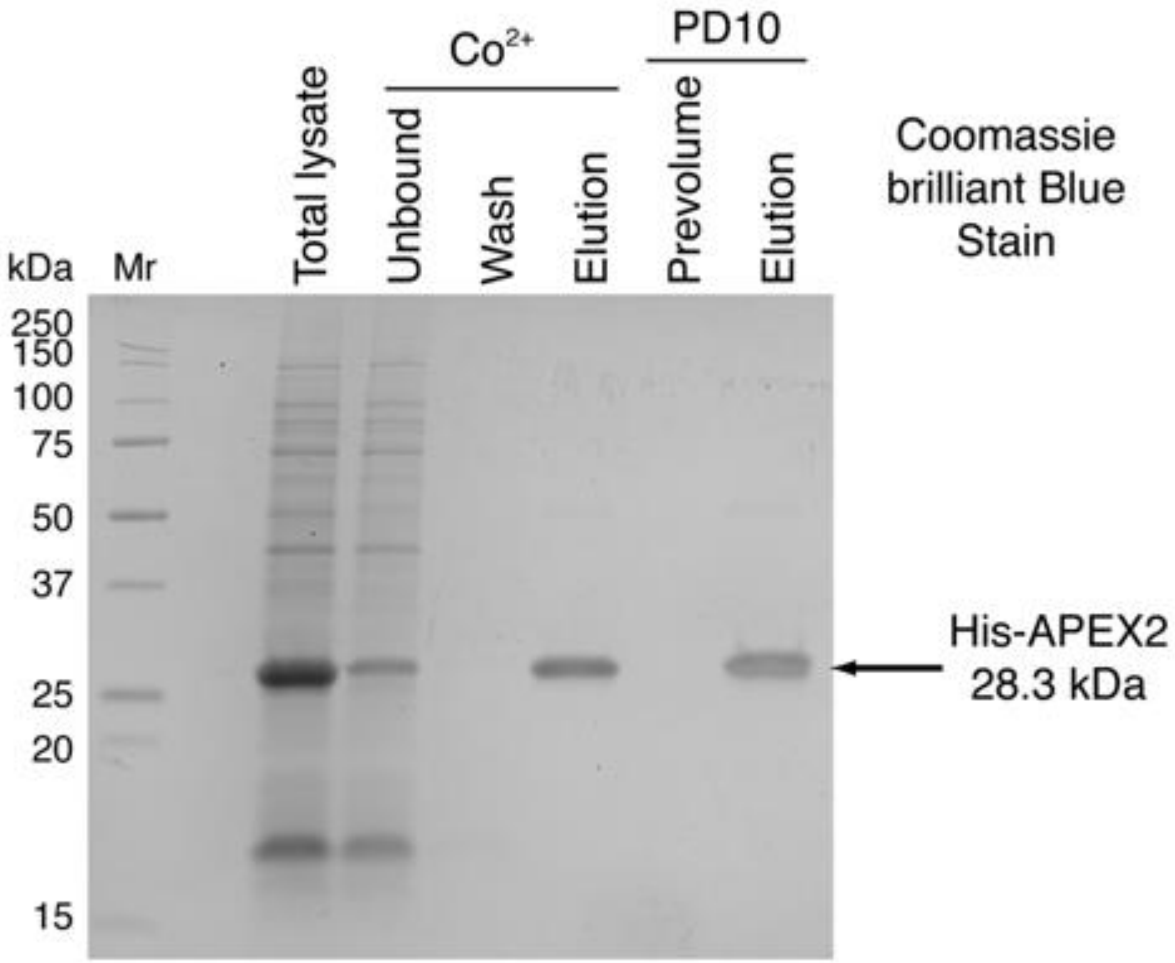


**Supplementary Figure S1.** SDS-PAGE analysis of the purification of soluble recombinant His-APEX2 protein. Starting from bacterial lysate, the purification workflow consists of cobalt-affinity chromatography, concentration using spin filter, buffer exchange with PD10 column, heme reconstitution, excess heme removal, and final spin filter concentration. The yields of APEX2 protein preparations are consistently >20 mg per 1 L of bacterial culture, and final densitometric purities >95%. Mr, relative molecular weight; kDa, kilodalton.

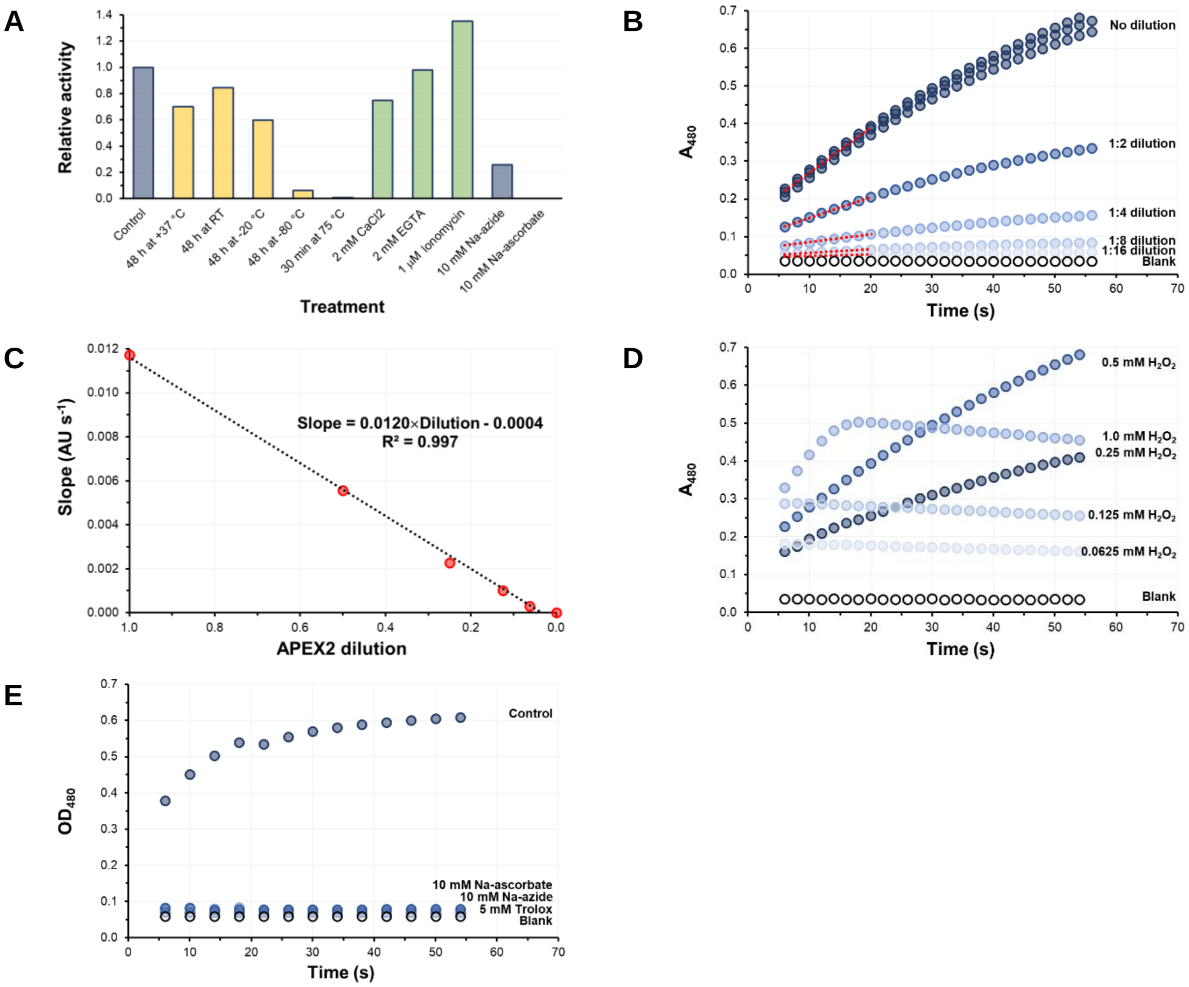


**Supplementary Figure S2.** Effects of storage conditions, the concentration of reagents, and the dilution of APEX2 to the peroxidase activity. (**A**) Effect of the enzyme storage conditions, additives, and quenchers on APEX2 activity. The presented activities are reported relative to untreated freshly purified APEX2 enzyme kept at +4 °C. The data indicate that freezing either at -20 or -80 °C will reduce or completely abolish the APEX2 activity, we thus stored purified APEX2 proteins at + 4 °C and freezing of APEX2 preparations was avoided altogether. (**B**) Absorbance versus time curves from the colorimetric activity assay at various APEX2 dilutions. The APEX2 activity is equal to the initial (linear) slope of the absorbance versus time curve (*red dashed lines*). The high reproducibility of the assay is illustrated by the low coefficient of variation (< 3%) of triplicate measurements of the highest dilution used. (**C**) The initial slope of the absorbance versus time curve of the colorimetric assay is linearly correlated with the used APEX2 dilution, indicating the assay's suitability for gauging and normalizing APEX2 activities. (**D**) Effect of $H_2O_2$ concentration on APEX2 activity. Inhibitory effect of $H_2O_2$ is visible at concentrations higher than 0.5 mM. (**E**) Optical density versus time curves of turbidimetric assay utilizing the formation of colloid, presumably from cross-linked BT and APEX2.

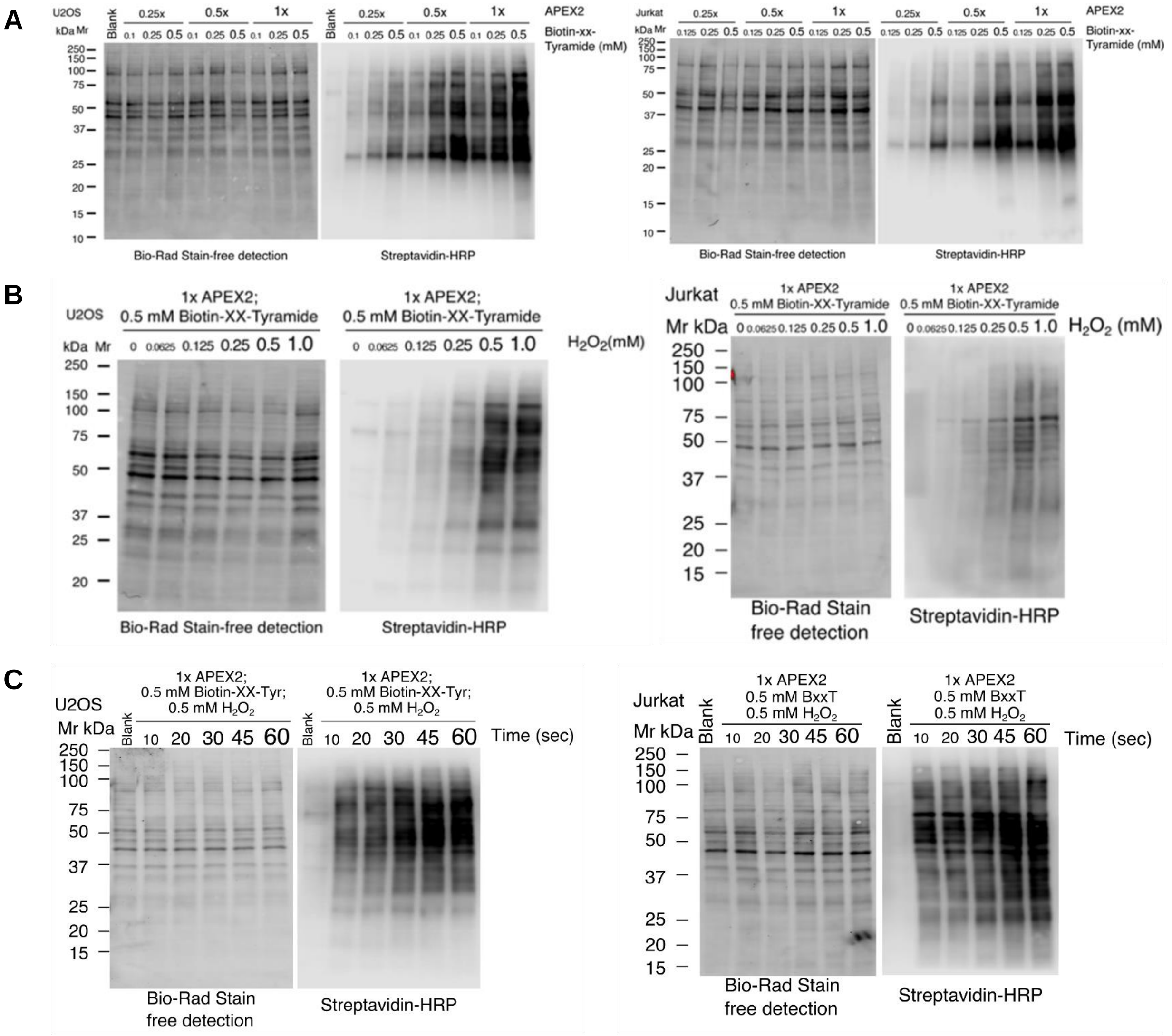


**Supplementary Figure S3.** Optimization of SLAPSHOT labeling conditions using adherent U2OS (*panel doublets on the left*) and suspended Jurkat cells (*panel doublets on the right*). The optimized parameters included (**A**) APEX2 activity and BxxT concentration, (**B**) $H_2O_2$ concentration, (**C**) Labeling duration at room temperature. Each experiment includes the detection of total protein on the blotting membrane by stain-free imaging (*left panels*) and biotin incorporation by Streptavidin-blotting (*right panels*). The data indicate that 30 or 45 s labeling respectively for adherent U2OS and suspended Jurkat cells using 0.0005 AU $s^{-1}$ $\mu L^{-1}$ (1× activity) (as measured by a colorimetric assay, see **Supplementary Figure S2**) and 0.5 mM BxxT and 0.5 mM $H_2O_2$ yield robust labeling. The $H_2O_2$ optimization corroborates the results obtained from Streptavidin-blotting **(Supplementary Figure S2D**). Mr, relative molecular weight; kDa, kilodalton.

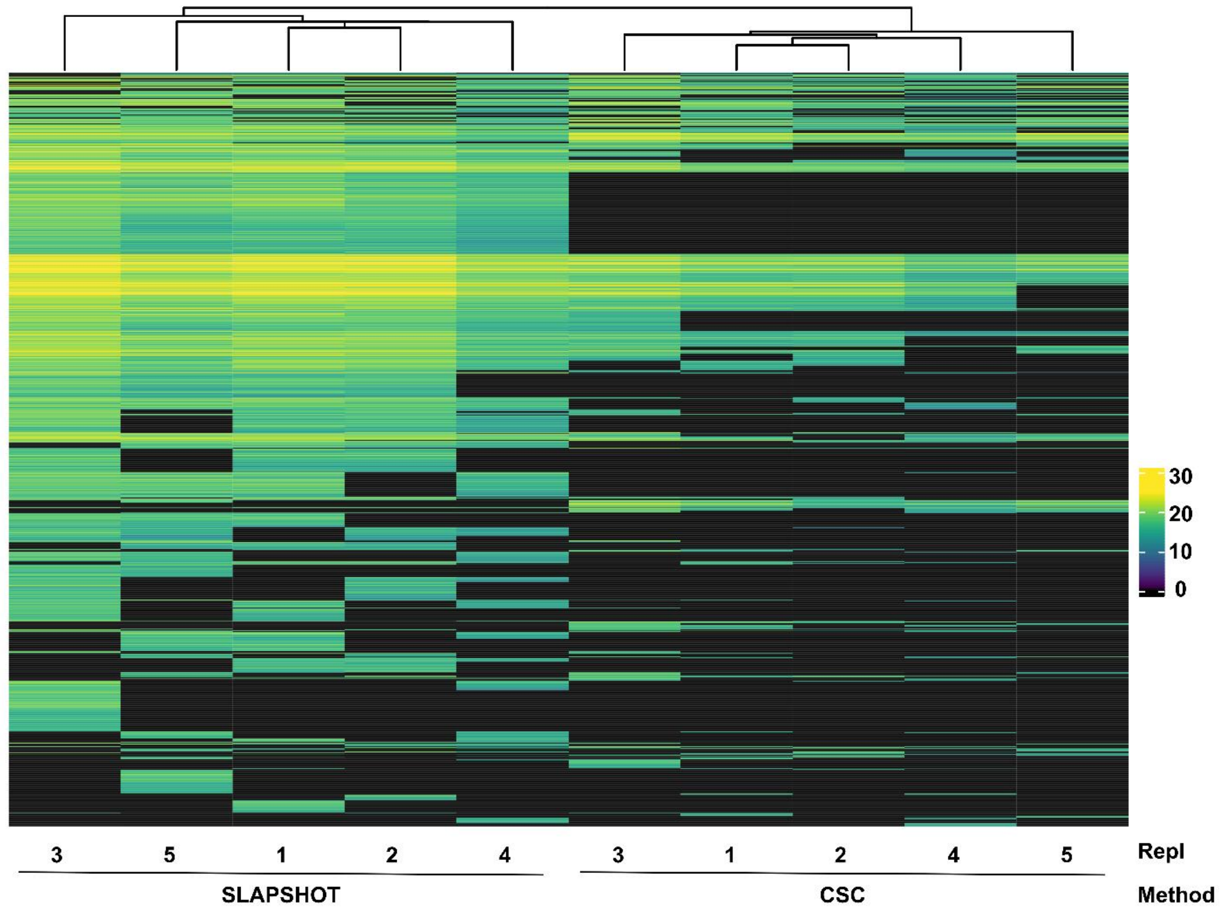


**Supplementary Figure S4.** Hierarchical clustering of riBAQ intensities from U2OS cells labeled with SLAPSHOT or CSC. The riBAQ intensities cluster most strongly based on the method used, indicating that the methods label different subset of proteins.

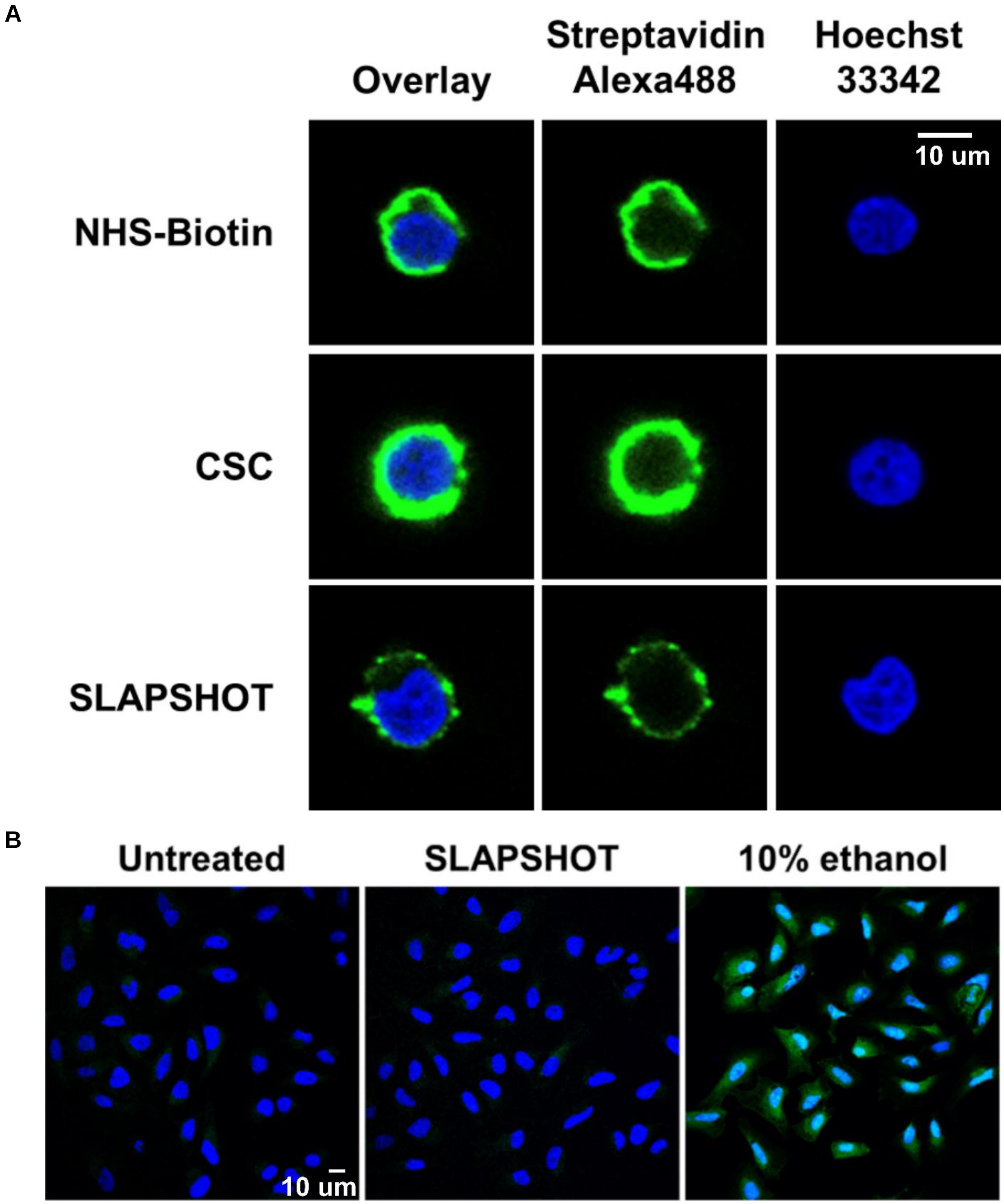


**Supplementary Figure S5.** SLAPSHOT preserves cellular integrity and selectively labels extracellularly exposed proteins. (**A**) Live-or-Dye assay on U2OS cells. The nuclei of all cells are labeled with Hoechst stain (*blue*), whereas Live-or-Dye reagent (*green*) only labels the cytoplasm of cells whose plasma membrane is compromised with 10% ethanol treatment. SLAPSHOT-treated cells are indistinguishable from untreated cells, indicating that SLAPSHOT does not compromise the integrity of plasma membrane. (**B**) Microscopic examination of Jurkat cells after biotinylating the cells with either NHS-Biotin, CSC or SLAPSHOT (all *green*). The nuclei of all cells are labeled with Hoechst stain (*blue*). The data indicate that the biotinylation is restricted to the cell periphery using all three methods.

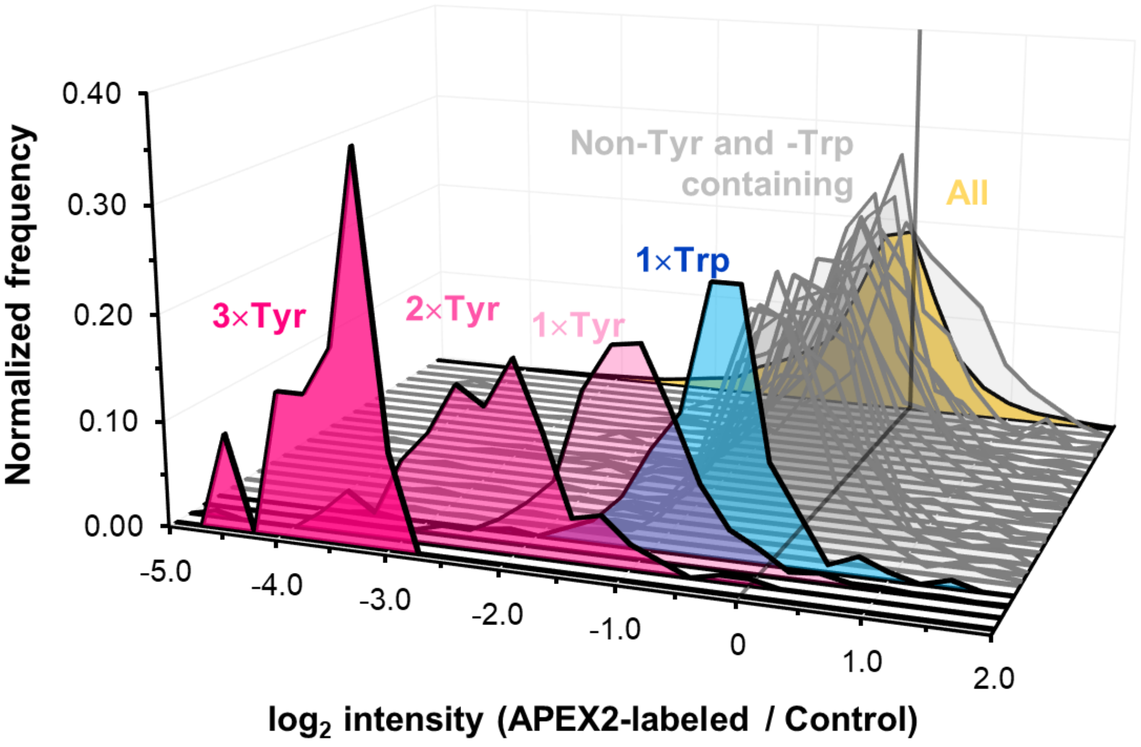


**Supplementary Figure S6.** Phenoxyl radicals generated by APEX2 target solely Tyr and Trp residues in proteins. Alkylated tryptic peptide mixture was treated with APEX2 in the presence of BP and $H_2O_2$, or left as an untreated control by omitting $H_2O_2$. After depletion of the biotinylated materials from both reactions, the two supernatants were individually analyzed by quantitative MS in data-dependent acquisition mode. $Log_2$ (APEX2-labeled/Control) intensity ratio was calculated for each peptide that was identified in both reactions. When considering all identified peptides ($N_{all}$ = 11731, in *yellow*), the distribution of log ratios centers near zero, indicating on average similar peptide intensities in both samples. As the analysis was performed label-free in order not to block any amino acid residues with the quantitation reagent, the distribution has significant width. Only Tyr (*pink*) and Trp (*blue*) containing peptides have log intensity ratio distributions that center below 0, indicating their depletion from the APEX2-labeled mixture compared to the control mixture ($N_{1\times Tyr}$ = 1603 $N_{2\times Tyr}$ = 171, $N_{3\times Tyr}$ = 22, $N_{1\times Trp}$ = 275). In these calculations, only those Tyr-containing peptides that do not contain Trp, and Trp-containing peptides that do not contain Tyr were included. Depletion efficiency increases linearly as the number of Tyr-residues in the peptide increases, as would be expected for phenoxyl radical targets. Due to the scarcity of Trp-containing residues, there were not enough peptides with two or more Trp residues for statistical analysis, but single Trp residue nevertheless exhibits depletion (appr. 40% of that of peptides with a single Tyr residue, indicating lesser reactivity). The other eighteen traces (*gray*) represent peptides that do not contain Tyr or Trp residues, but contain multiple residues of certain type (*e.g.*, three Ala, $N_{3\times\mathrm{Ala}}$ = 2471). Distributions of the log intensity ratios of these peptides center at zero, indicating that peptides containing even as many as three Ala residues are not effectively depleted from the biotinylated peptide mixture and thus Ala is not a target for the phenoxyl radicals. The Cys sulfhydryls were alkylated with iodoacetamide during peptide generation prior to APEX2 labeling, thus resembling sulfhydryls in most extracellularly exposed Cys residues that are blocked by disulfide bonding.

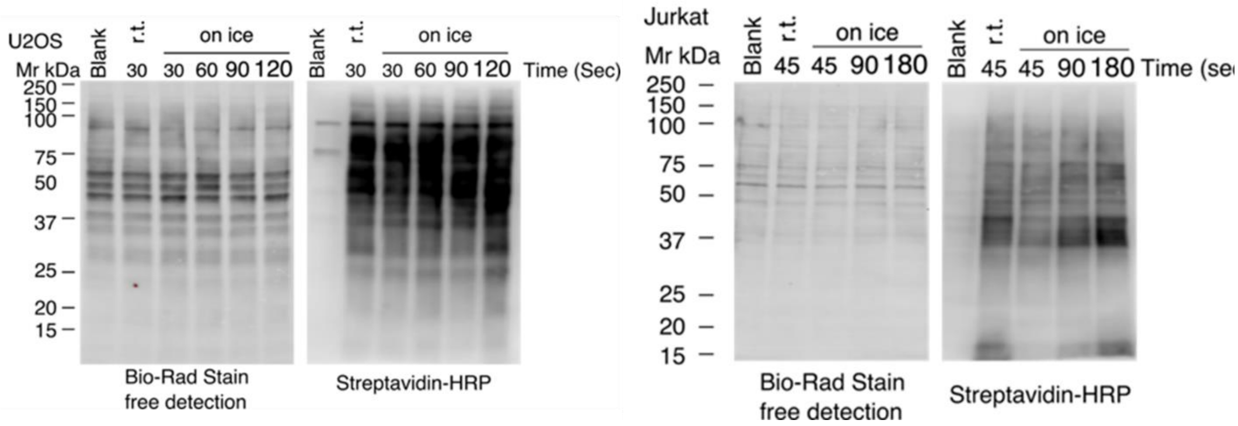


**Supplementary Figure S7.** Optimization of SLAPSHOT labeling duration of adherent U2OS (*panel doublet on the left*) and suspended Jurkat (*panel doublet on the right*) cells on ice. Each experiment includes the detection of total protein on the blotting membrane by stain-free imaging (*left panels*) and biotin incorporation by Streptavidin-blotting (*right panels*). The data indicate that doubling the labeling time used for room temperature labeling, to 60 and 90 s for adherent U2OS and suspended Jurkat cells yields robust labeling. Mr, relative molecular weight; kDa, kilodalton.

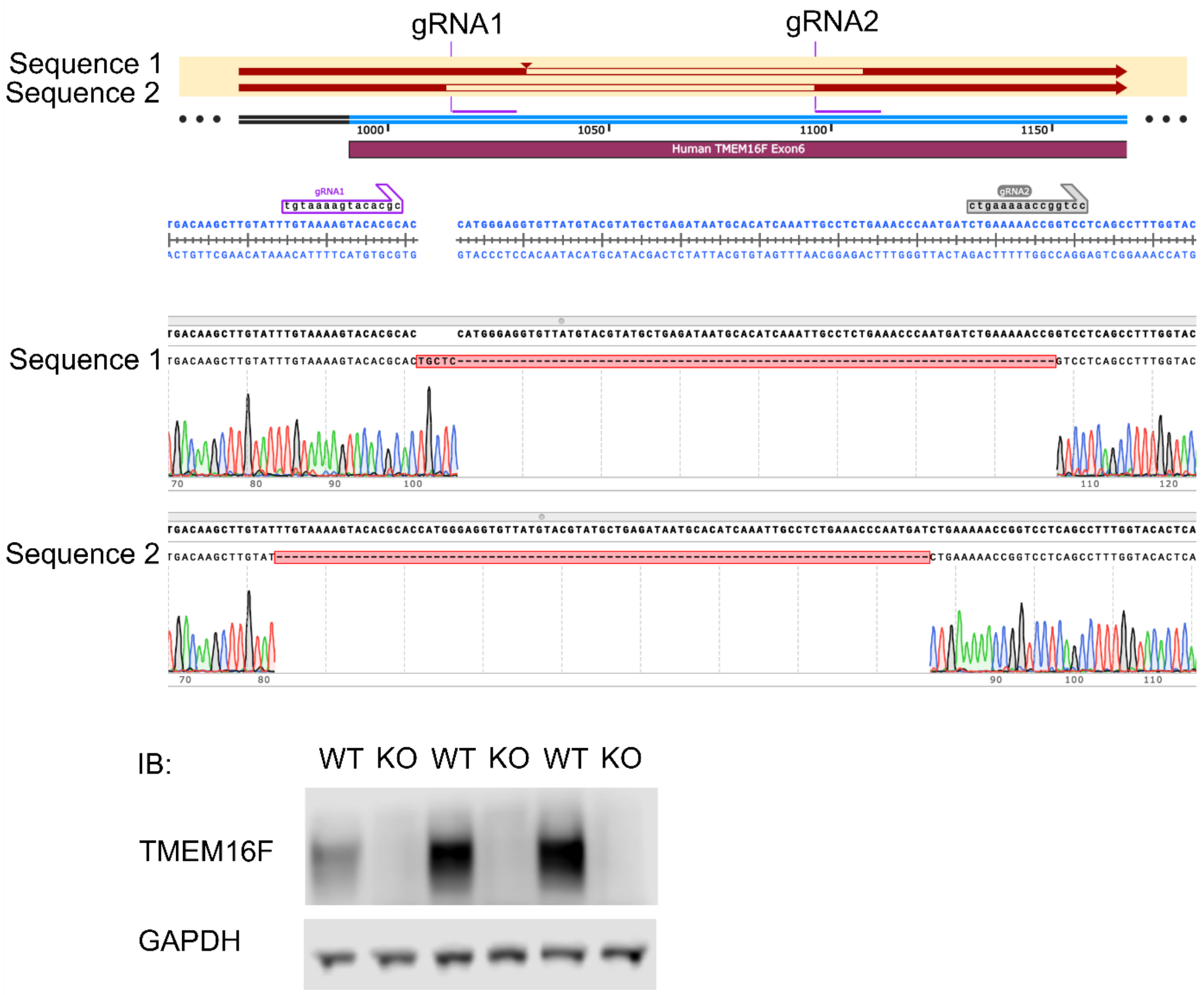


**Supplementary Figure S8.** Genetic and biochemical verification of *ANO6* CRISPR-Cas9 knock-out in Jurkat cells. Sanger sequencing results indicate that the cell population is either a mixture of two clones both with homozygous deletions in the exon 6 of *ANO6*, or a single heterozygous clone. The deletions lead to frameshifts and premature stop codons that create a truncated transcript and undetectable TMEM16F protein. Western blotting of three individually expanded clones of both WT and TMEM16F KO cells indicates absence of TMEM16F protein in the knock-out cells.

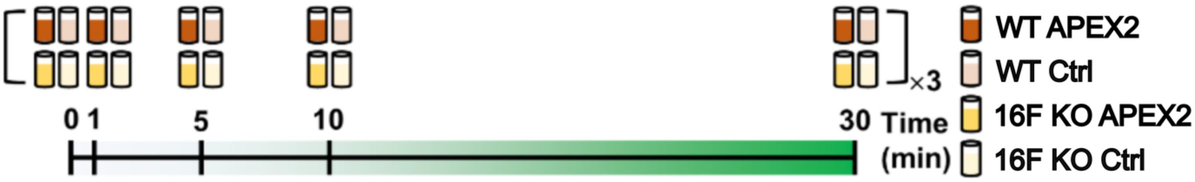


**Supplementary Figure S9.** Schematic of the time-course ionomycin stimulation of Jurkat cells. WT and 16F KO cells were stimulated with bacterial calcium ionophore ionomycin for durations ranging from 1 to 30 min. Cells without stimulation (0 min, or pre-stimulation) were included as controls. The samples were labeled with SLAPSHOT according to the optimized protocol. Matching control without $H_2O_2$ was included for each individual labeled sample. The experiment was replicated three times using different passages of the cells.

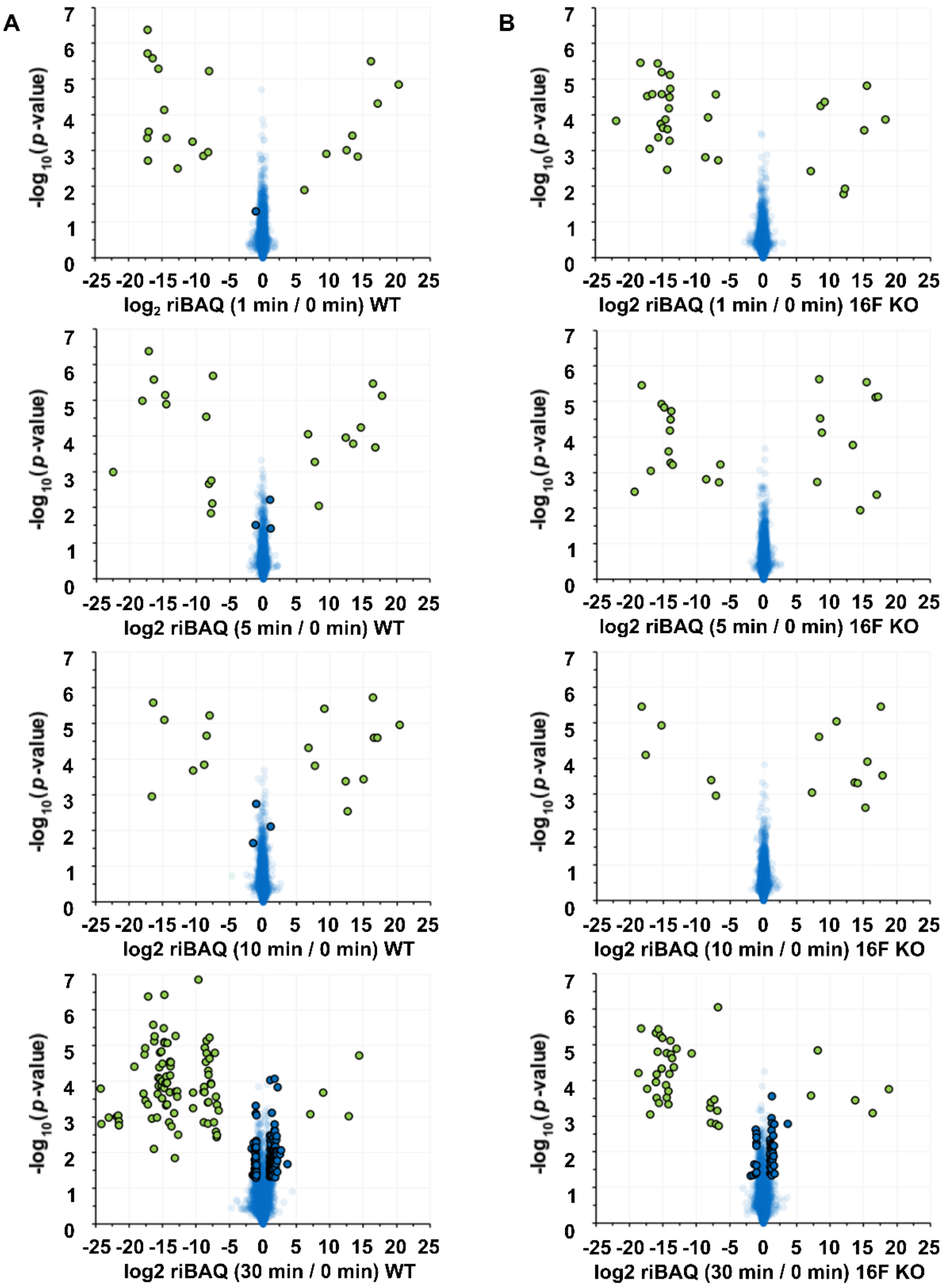


**Supplementary Figure S10.** Volcano plots indicating the fold-change and significance of protein intensities between variously ionomycin treated (1, 5, 10 or 30 min) (**A**) WT and (**B**) 16F KO Jurkat cells and non-treated (0 min) controls. Datapoints in *blue* indicate proteins that were detected in both samples, whereas datapoints in *green* indicate proteins that were identified in just one sample and imputed in the other (the *green* datapoints are included merely for visualization purposes). Proteins whose absolute $\log_2$ riBAQ fold-change >1, and -$\log_{10}$ *p*-value >1.3 (*p*-value < 0.05) are *circled*. The data indicate that relatively small number of proteins are differentially detected between 0 min and either 1, 5, or 10 min stimulation time-points in both WT and 16F KO cells, whereas larger number of proteins are differentially detected between 0 min and 30 min stimulations in both cell types.

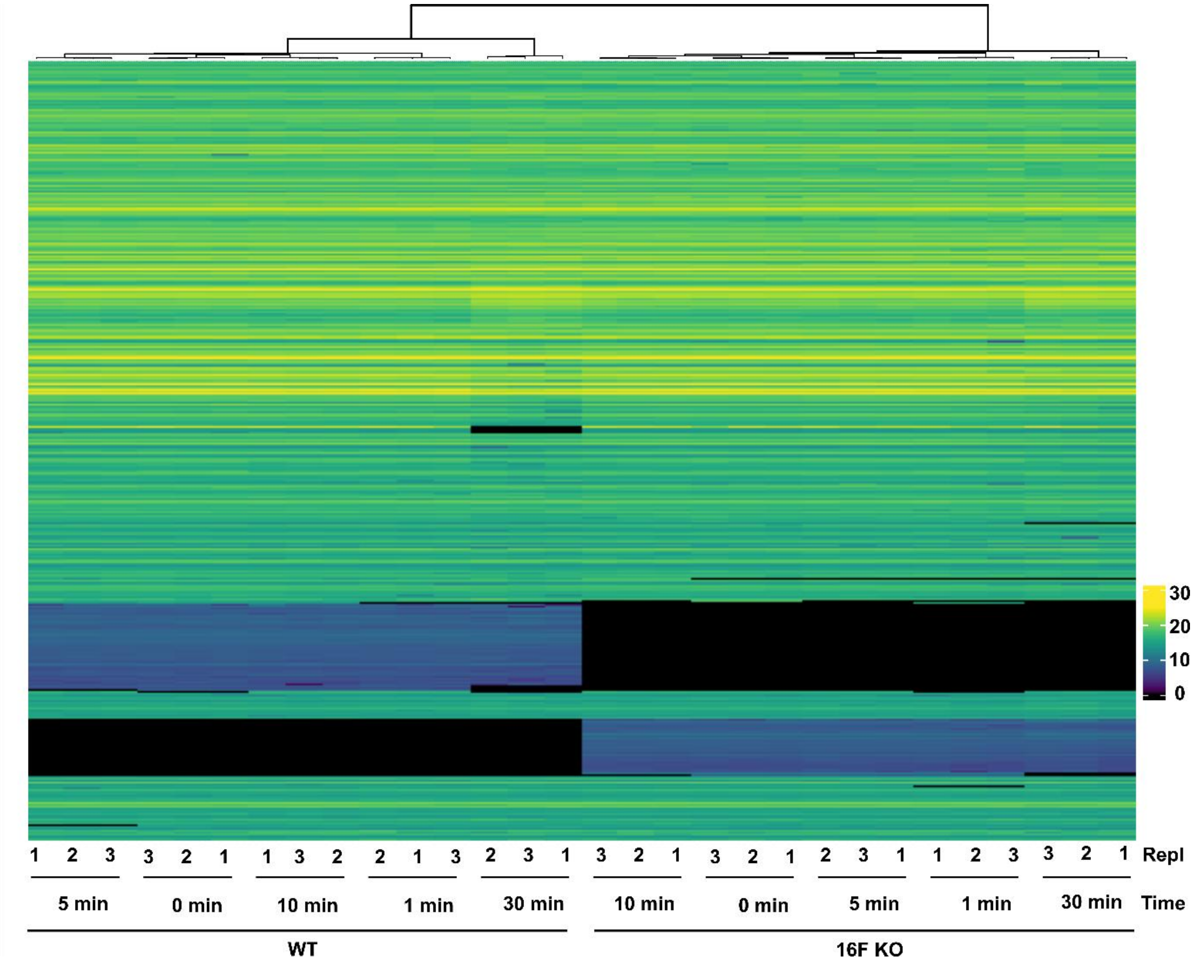


**Supplementary Figure S11.** Hierarchical clustering of WT and 16F KO Jurkat cell ionomycin stimulation time-course data. The TMT-riBAQ intensities separate strongly based on the cell type, followed by time point. The data indicates strong reproducibility of the triplicate measurements from a given cell type and time point.

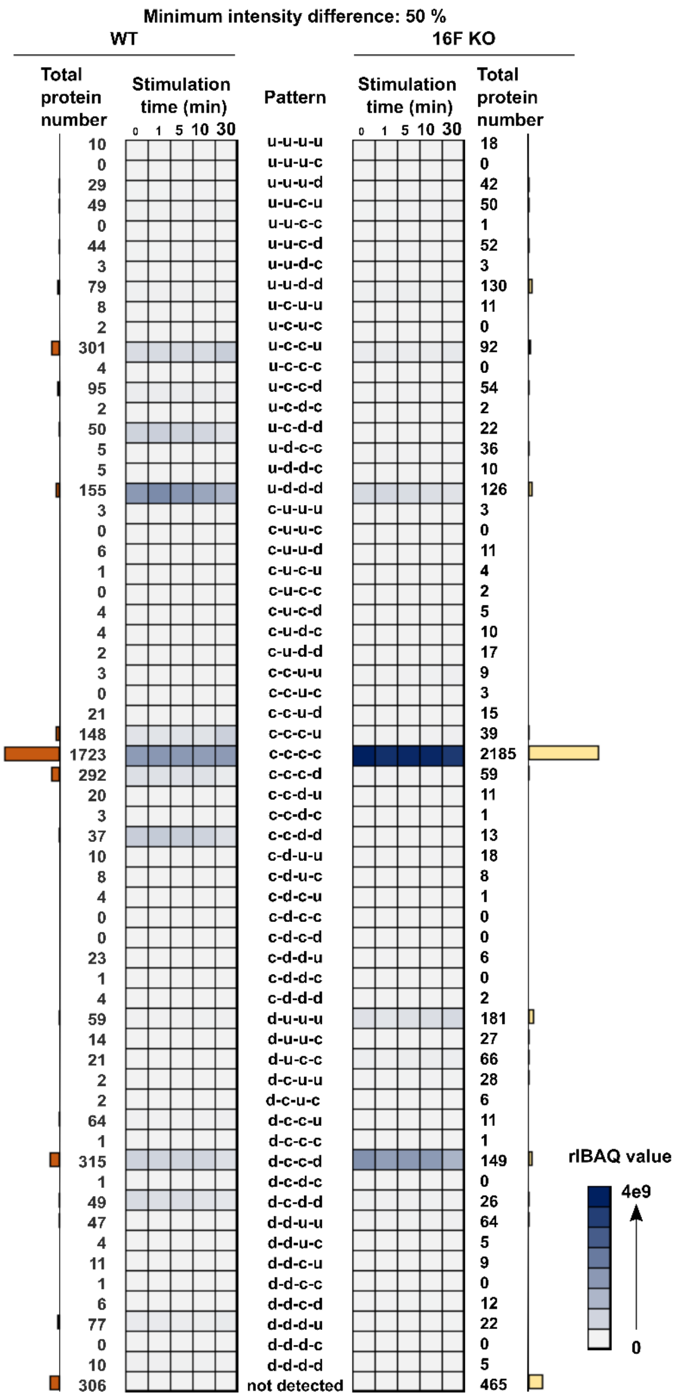


**Supplementary Figure S12.** Heatmaps showing the evolution of the total riBAQ for intensity patterns in WT (*left heatmap*) and 16F KO (*right heatmap*) cells, when considering all detected proteins ($N$ = 4148). Each square in the grid represents the sum of riBAQ values from the proteins with that intensity pattern. The total number of proteins belonging to a pattern is shown by bar graph.  Notably, most of the proteins as well as riBAQ intensity is concentrated in a few patterns.

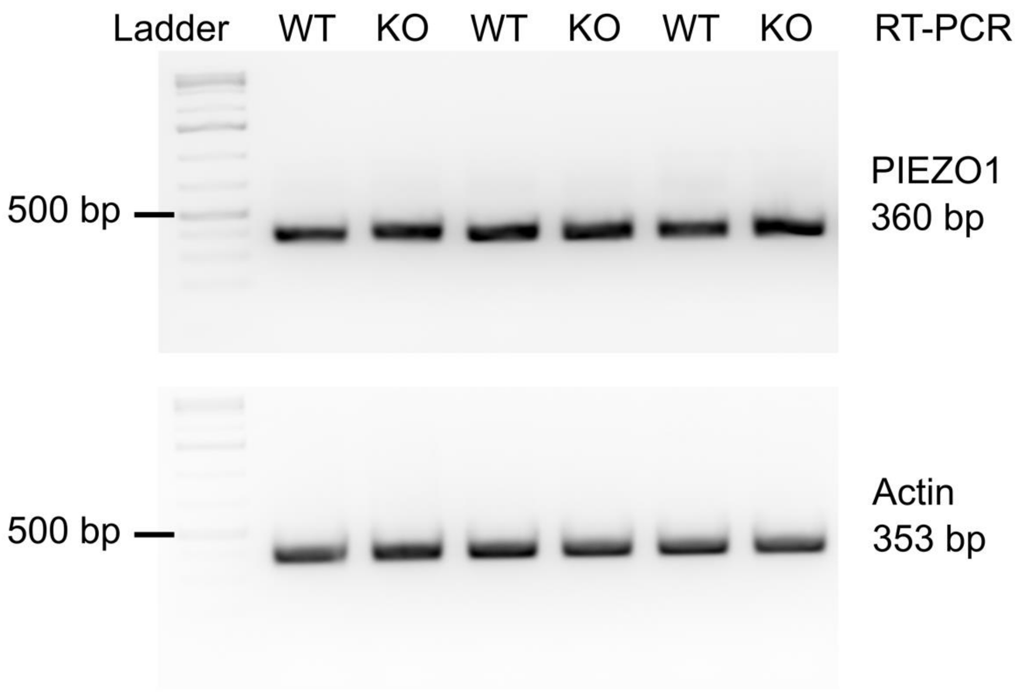


**Supplementary Figure S13.** RT-PCR products of PIEZO1 transcript (*upper panel*) in WT and 16F KO Jurkat cells with actin transcript (*lower panel*) as a control. Data of three replicates of both WT and TMEM16F KO cells indicates absence of PIEZO1 transcript in the knock-out cells. bp, base pairs.

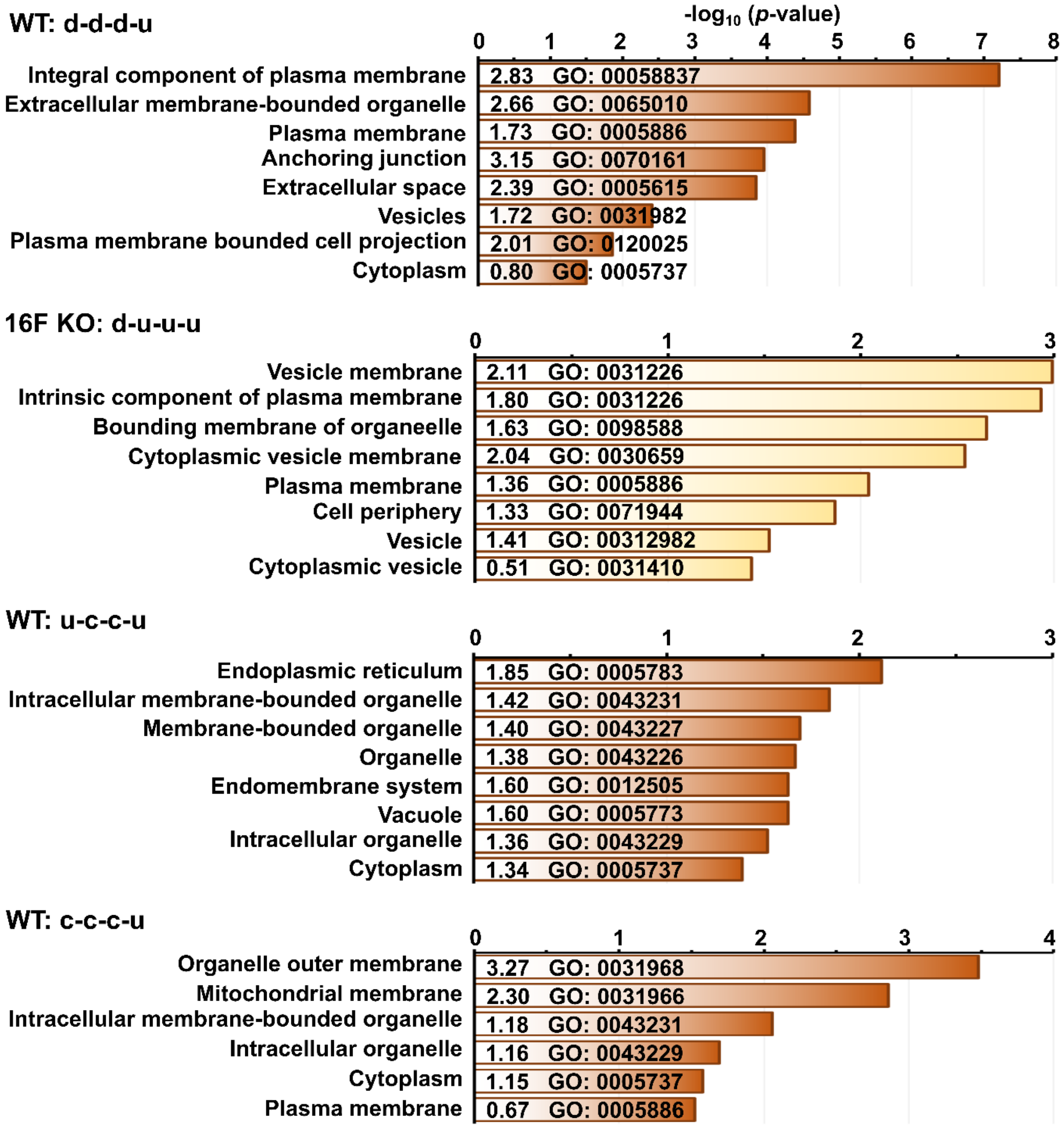


**Supplementary Figure S14.** PANTHER analysis (GO cellular component) of proteins with selected intensity patterns. Length of the bar indicates significance with $-\log_{10}$ (*p*-value). GO terms and descriptions are shown, as are the enrichment factors. Enrichment factors > 1 indicate overrepresentation of proteins in that category in the data, whereas enrichment factor < 1 indicates depletion.

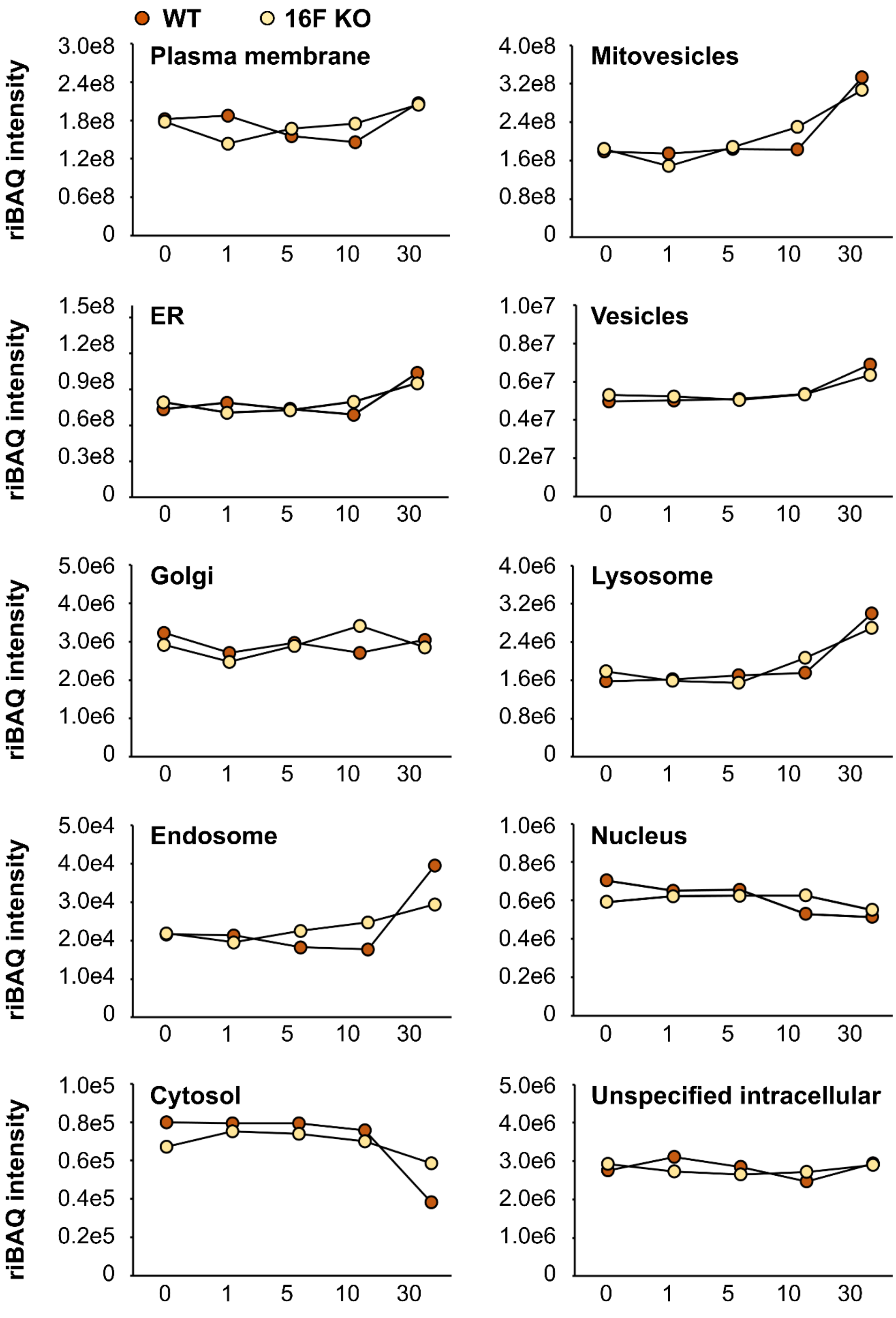


**Supplementary Figure S15.** Total riBAQ intensity patterns of proteins with a selected localization annotation.

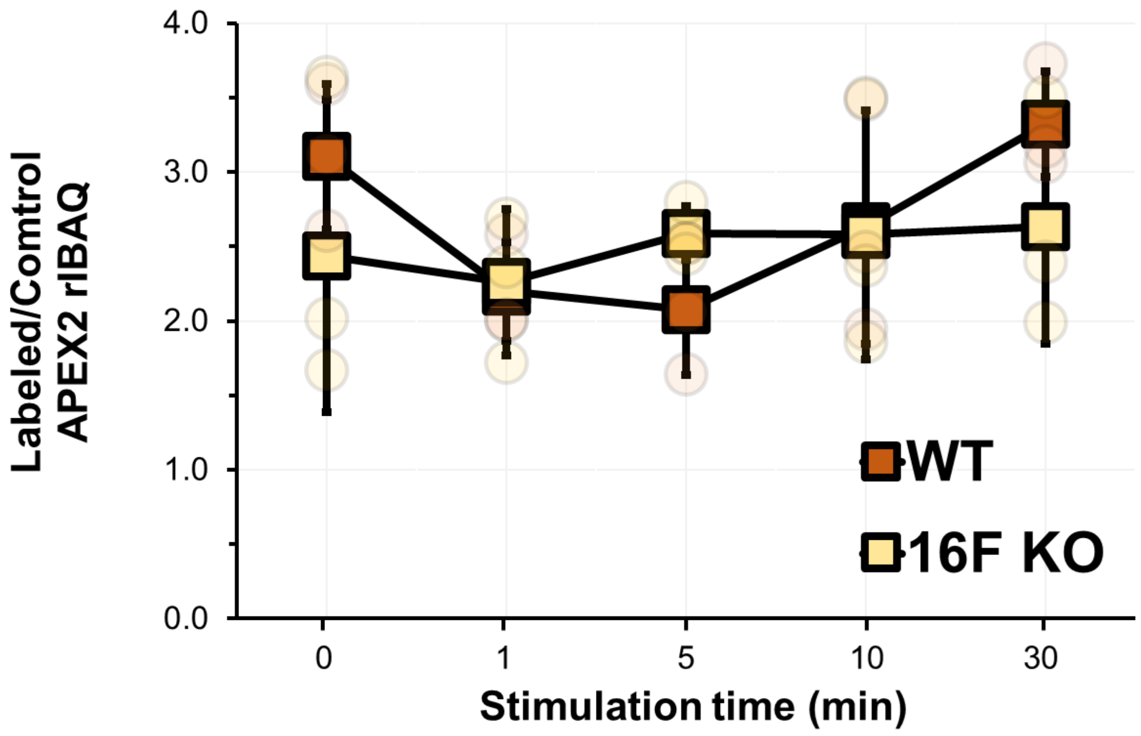


**Supplementary Figure S16.** His-APEX2 protein riBAQ ratios from the labeled and control samples, with mean ± standard deviation indicated, *N* = 3. The data indicate that cell pellets from the labeling reactions contain 2.5-fold more APEX2 protein compared to the non-labeled control cell pellets. This low and very consistent ratio indicates that there is no major cellular internalization of the His-APEX2 protein during the labeling reaction, either by calcium-dependent (*e.g.*, endocytosis) or other processes. Rather, non-specific binding to cells, plasticware and Neutravidin-agarose contribute to the His-APEX2 riBAQ intensity in the samples.

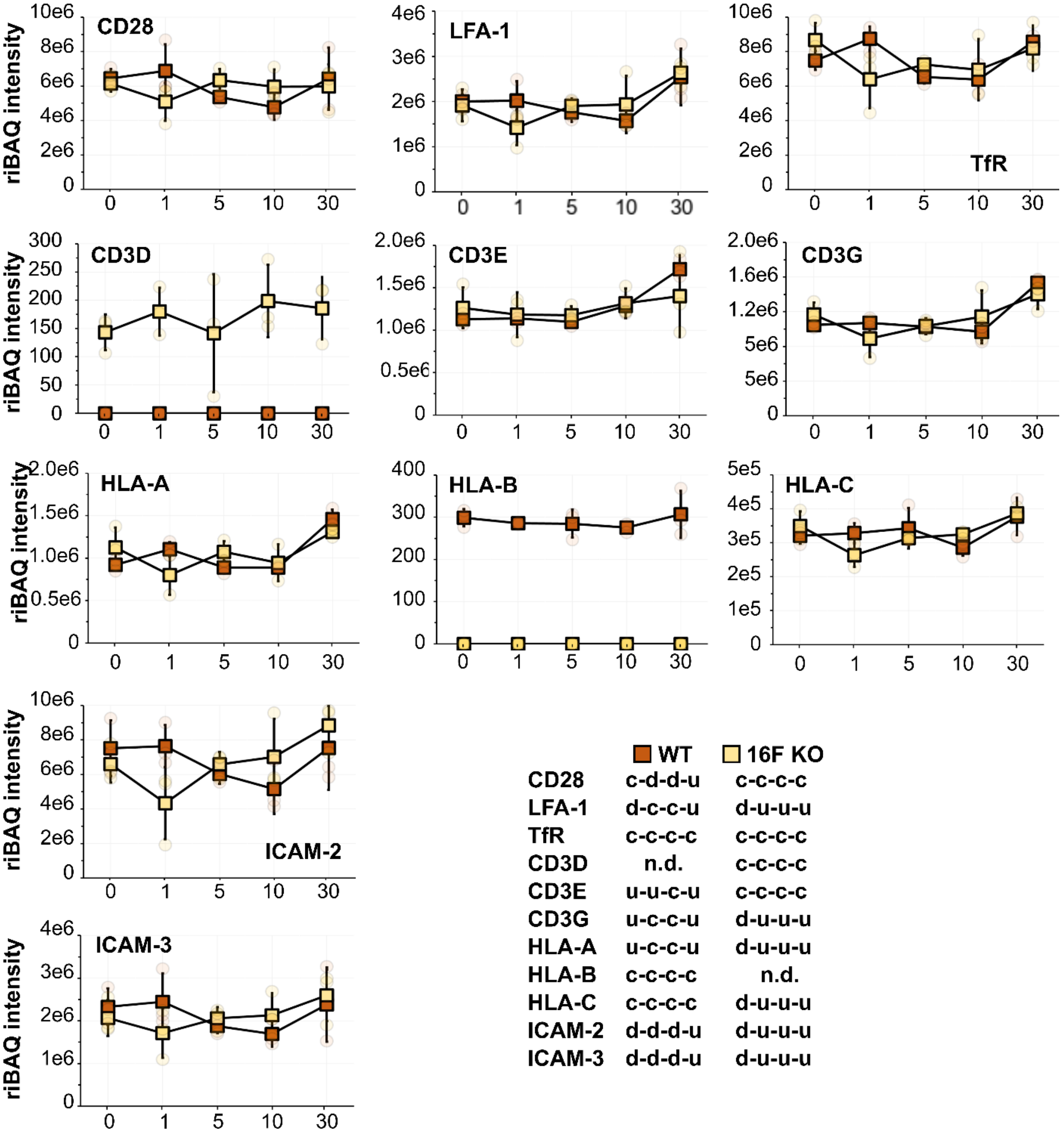


**Supplementary Figure S17.** MS time-course profiles of proteins that were monitored by Bricogne et al., 2019 using PD-1 overexpressing Jurkat cells with 15 min of 5 µM ionomycin stimulation, or proteins in the same family. n.d., not detected.
