## Supplementary R code for "SLAPSHOT reveals rapid dynamics of extracellularly exposed proteome in response to calcium-activated plasma membrane phospholipid scrambling"

## ----------------------------------------------------------------------------------
##
## SLAPSHOT reveals rapid dynamics of extracellularly exposed proteome in response to calcium-activated plasma membrane phospholipid scrambling
## Sami T. Tuomivaara*, Chin Fen Teo*, Yuh Nung Jan, Lily Y. Jan#, Arun P. Wiita#
##
## Supplementary R code
##
##
## These code snippets allow reproduction of the analyses described in the title manuscript. 
##
## ----------------------------------------------------------------------------------


## ----------------------------------------------------------------------------------
##
## Data imputation
##
## ----------------------------------------------------------------------------------

## Some help at https://github.com/farrellday/miceRanger

library(miceRanger)
library(doParallel)

## Open a tab-separated file with the following structure:
## Protein IDs in column 1
## The corresponding background-corrected log2-transformed intensities from both WT and 16F KO cells
## in columns 2 to 31
## NA entries will be imputed

miceRanger.in <- read.table(file.choose(),
                            header = TRUE,
                            sep = "\t",
                            row.names = 1)

cl <- makeCluster(detectCores(logical = TRUE) - 1)
registerDoParallel(cl)

miceRanger.out <- miceRanger(miceRanger.in,
                             m = 10,
                             returnModels = FALSE,
                             verbose = TRUE,
                             maxiter = 25,
                             parallel = TRUE,
                             num.trees = 100)

stopCluster(cl)
registerDoSEQ()

plotDistributions(miceRanger.out, vars = 'allNumeric')
plotCorrelations(miceRanger.out, vars = 'allNumeric')
plotVarConvergence(miceRanger.out, vars = 'allNumeric')
plotModelError(miceRanger.out, vars = 'allNumeric')
plotVarImportance(miceRanger.out)
plotImputationVariance(miceRanger.out, ncol = 2, widths = c(5, 3))

final.miceRanger.out <- miceRanger.in
final.miceRanger.out[is.na(miceRanger.in)] <- 1

for (i in 1:length(miceRanger.out$finalImps))
{
  for (j in 1:ncol(final.miceRanger.out))
  {
    imputed <- which(miceRanger.out$naWhere[, j])
    values <- miceRanger.out$finalImps[[i]][[j]]
    final.miceRanger.out[imputed, j] <- final.miceRanger.out[imputed, j] * values^(1 / length(miceRanger.out$finalImps))
  }
}

final.miceRanger.out <- 2^final.miceRanger.out


## ----------------------------------------------------------------------------------
##
## Internal Reference Scaling (IRS) with pseudo-reference
##
## ----------------------------------------------------------------------------------

## Some help at https://pwilmart.github.io/IRS_normalization/understanding_IRS.html

library(edgeR)

## Open a tab-separated file with the following structure:
## Protein IDs in column 1
## The corresponding (not log2-transformed) intensities from both WT and 16F KO cells
## in columns 2 to 31

irs.in <- read.table(file.choose(),
                     header = TRUE,
                     row.names = 1,
                     sep = "\t")

irs.in[irs.in == 0] <- 1
irs.in <- as.data.frame(irs.in)

irs.exp1.raw <- irs.in[1:5]
irs.exp2.raw <- irs.in[6:10]
irs.exp3.raw <- irs.in[11:15]
irs.exp4.raw <- irs.in[16:20]
irs.exp5.raw <- irs.in[21:25]
irs.exp6.raw <- irs.in[26:30]

target <- mean(c(colSums(irs.exp1.raw),
                 colSums(irs.exp2.raw),
                 colSums(irs.exp3.raw),
                 colSums(irs.exp4.raw),
                 colSums(irs.exp5.raw),
                 colSums(irs.exp6.raw)))

norm_facs <- target / colSums(irs.exp1.raw)
irs.exp1.sl <- sweep(irs.exp1.raw, 2, norm_facs, FUN = "*")

norm_facs <- target / colSums(irs.exp2.raw)
irs.exp2.sl <- sweep(irs.exp2.raw, 2, norm_facs, FUN = "*")

norm_facs <- target / colSums(irs.exp3.raw)
irs.exp3.sl <- sweep(irs.exp3.raw, 2, norm_facs, FUN = "*")

norm_facs <- target / colSums(irs.exp4.raw)
irs.exp4.sl <- sweep(irs.exp4.raw, 2, norm_facs, FUN = "*")

norm_facs <- target / colSums(irs.exp5.raw)
irs.exp5.sl <- sweep(irs.exp5.raw, 2, norm_facs, FUN = "*")

norm_facs <- target / colSums(irs.exp6.raw)
irs.exp6.sl <- sweep(irs.exp6.raw, 2, norm_facs, FUN = "*")

irs <- data.frame(rowSums(irs.exp1.sl),
              rowSums(irs.exp2.sl),
              rowSums(irs.exp3.sl),
              rowSums(irs.exp4.sl),
              rowSums(irs.exp5.sl),
              rowSums(irs.exp6.sl))

colnames(irs) <- c("sum1", "sum2", "sum3", "sum4", "sum5", "sum6")

irs$average <- apply(irs, 1, function(x) exp(mean(log(x))))

irs$fac1 <- irs$average / irs$sum1
irs$fac2 <- irs$average / irs$sum2
irs$fac3 <- irs$average / irs$sum3
irs$fac4 <- irs$average / irs$sum4
irs$fac5 <- irs$average / irs$sum5
irs$fac6 <- irs$average / irs$sum6

data.irs <- irs.exp1.sl * irs$fac1
data.irs <- cbind(data.irs, irs.exp2.sl * irs$fac2)
data.irs <- cbind(data.irs, irs.exp3.sl * irs$fac3)
data.irs <- cbind(data.irs, irs.exp4.sl * irs$fac4)
data.irs <- cbind(data.irs, irs.exp5.sl * irs$fac5)
data.irs <- cbind(data.irs, irs.exp6.sl * irs$fac6)

irs_tmm <- calcNormFactors(data.irs)
irs.out <- sweep(data.irs, 2, irs_tmm, FUN = "/") 


## ----------------------------------------------------------------------------------
##
## Calculating distances between protein intensity profiles and theoretical models
##
## ----------------------------------------------------------------------------------

## Some help at https://cran.r-project.org/web/packages/philentropy/vignettes/Distances.html

library(philentropy)
library(foreach)
library(doParallel)

## Open a tab-separated file with the following structure:
## Model IDs in column 1
## Corresponding model intensities for five time-points in columns 2 to 6

profile.models <- read.table(file.choose(),
                             header = TRUE,
                             row.names = 1,
                             sep = "\t")

profile.models <- sweep(profile.models,
                        1,
                        rowSums(profile.models),
                        FUN = "/")

## Open a tab-separated file with the following structure:
## Protein IDs in column 1
## Corresponding empirical protein intensities from five time-points (each average from the triplicate measurements)
## from either WT or 16F KO cells in columns 2 to 6

profile.data <- read.table(file.choose(),
                           header = TRUE,
                           row.names = NULL,
                           sep = "\t")

profile.results <- data.frame(ID = profile.data[, 1], best.model = "not.detected")

profile.data <- profile.data[, -1]
colnames(profile.data) <- colnames(profile.models)
rows <- as.logical(rowSums(profile.data != 0))
profile.data[rows, ] <- profile.data[rows, ] / rowSums(profile.data[rows, ])

my.cluster <- parallel::makeCluster(20, type = "PSOCK")
doParallel::registerDoParallel(cl = my.cluster)

dists <- rep(0, length = nrow(profile.models))

for (i in 1:nrow(profile.data))
{
  cat(i, "of", nrow(profile.data), "\n")
  if (sum(profile.data[i, ]) != 0)
  {
    dists <- foreach(j = 1:nrow(profile.models), .export = "distance") %dopar% {
      distance(rbind(profile.models[j, ], profile.data[i, ]), method = "bhattacharyya", unit = "log", mute.message = TRUE)
                                                                               }
    profile.results[i, "best.model"] <- rownames(profile.models)[which.min(dists)]
  }
}

parallel::stopCluster(cl = my.cluster)


## ----------------------------------------------------------------------------------
##
## Fisher’s exact test for calculating p-values for under- and over-representation
##
## ----------------------------------------------------------------------------------

## Open a tab-separated file with the following structure:
## Matrix where each row denotes one of the 61 WT patterns, and each column denotes one of the 61 16F KO patterns
## Each element within the matrix, at the intersection of a given row and column
## denotes the number of proteins that have a particular pattern in the WT cells (indicated by the row)
## and a particular pattern in the 16F KO cells (indicated by the column)

hypergeomdata <- read.table(file.choose(),
                            header = FALSE,
                            row.names = NULL,
                            sep = "\t")

hypergeomdata <- as.matrix(hypergeomdata)

## Baseline (expected) overlap based on hypergeometric distribution assuming randomness
hypergeom.expected <-  matrix(0,
                              ncol = ncol(hypergeomdata),
                              nrow = nrow(hypergeomdata))

colnames(hypergeom.expected) <- colnames(hypergeomdata)
rownames(hypergeom.expected) <- rownames(hypergeomdata)

for(i in 1:nrow(hypergeom.expected)) for (j in 1:ncol(hypergeom.expected))
{
  if(sum(hypergeomdata[i, ]) <= 0 | sum(hypergeomdata[, j]) <= 0) hypergeom.expected[i, j] <- NA
  else
  {
    vec <- dhyper(0:sum(hypergeomdata[i, ]), sum(hypergeomdata[, j]), sum(hypergeomdata) - sum(hypergeomdata[, j]), sum(hypergeomdata[i, ]))
    hypergeom.expected[i, j] <- which(vec == max(vec)) - 1
  }
}

## Enrichment factor
hypergeom.enrichment <- matrix(0,
                               ncol = ncol(hypergeomdata),
                               nrow = nrow(hypergeomdata))

colnames(hypergeom.enrichment) <- colnames(hypergeomdata)
rownames(hypergeom.enrichment) <- rownames(hypergeomdata)

for (i in 1:nrow(hypergeom.enrichment)) for (j in 1:ncol(hypergeom.enrichment))
{
  if(is.na(hypergeom.expected[i, j])) hypergeom.enrichment[i, j] <- NA
  else hypergeom.enrichment[i, j] <- (hypergeomdata[i, j] * sum(hypergeomdata)) / (sum(hypergeomdata[i, ]) * sum(hypergeomdata[, j]))
}

## Fisher's exact test for over-representation
hypergeom.over <- matrix(0,
                         ncol = ncol(hypergeomdata),
                         nrow = nrow(hypergeomdata))

colnames(hypergeom.over) <- colnames(hypergeomdata)
rownames(hypergeom.over) <- rownames(hypergeomdata)

for(i in 1:nrow(hypergeom.over)) for (j in 1:ncol(hypergeom.over))
{
  hypergeom.over[i, j] <- phyper(hypergeomdata[i, j] - 1,
                                 sum(hypergeomdata[, j]),
                                 sum(hypergeomdata) - sum(hypergeomdata[, j]),
                                 sum(hypergeomdata[i, ]),
                                 lower.tail = FALSE)
}

## Fisher's exact test for under-representation
hypergeom.under <- matrix(0,
                          ncol = ncol(hypergeomdata),
                          nrow = nrow(hypergeomdata))

colnames(hypergeom.under) <- colnames(hypergeomdata)
rownames(hypergeom.under) <- rownames(hypergeomdata)

for(i in 1:nrow(hypergeom.under)) for (j in 1:ncol(hypergeom.under))
{
  hypergeom.under[i, j] <- phyper(hypergeomdata[i, j],
                                  sum(hypergeomdata[, j]),
                                  sum(hypergeomdata) - sum(hypergeomdata[, j]),
                                  sum(hypergeomdata[i, ]),
                                  lower.tail = TRUE)
}


## ----------------------------------------------------------------------------------
##
## Hierarchical clustering
##
## Supplementary Figures S4 and S11
##
## ----------------------------------------------------------------------------------

library(ComplexHeatmap)
library(circlize)
library(viridis)
library(svglite)

## Open a tab-separated file with the following structure:
## Each column corresponds to MS intensities of a sample, no ID column

hierclust.in <- read.table(file.choose(),
                           header = TRUE,
                           row.names = 1,
                           sep = "\t",
                           stringsAsFactors = FALSE)

hierclust.in <- as.matrix(hierclust.in)

col_fun = colorRamp2(c(0, 1:25),
                     c("black", viridis(25)))

dev.new(width = 1, height = 1, unit = "in")
Heatmap(hierclust.in,
        clustering_distance_rows = "pearson",
        clustering_distance_columns = "pearson",
        clustering_method_rows = "single",
        clustering_method_columns = "single",
        na_col = "black",
        col = col_fun)

#This script saves the file on disk
svglite("clustering.svg", width = 1, height = 1)

plot(Heatmap(hierclust.in,
             clustering_distance_rows = "spearman",
             clustering_distance_columns = "spearman",
             clustering_method_rows = "complete",
             clustering_method_columns = "complete",
             na_col = "black",
             col = col_fun))
dev.off()


## ----------------------------------------------------------------------------------
##
## Sankey flow diagram
##
## Figure 5
##
## ----------------------------------------------------------------------------------

## Some help at https://christophergandrud.github.io/networkD3/

library(networkD3)

nodes = data.frame("name" = c("WT u-c-c-u",
                              "WT u-d-d-d",
                              "WT c-c-c-u",
                              "WT c-c-c-c",
                              "WT d-d-d-u",
                              "WT not.detcted",
                              "16F KO u-c-c-u",
                              "16F KO c-c-c-u",
                              "16F KO c-c-c-c",
                              "16F KO d-u-u-u",
                              "16F KO d-c-u-u",
                              "16F KO d-d-u-u",
                              "16F KO not.detected"))

nodes$nodegroups <- c("gr1","gr1","gr1","gr1","gr1","gr1","gr2","gr2","gr2","gr2","gr2","gr2", "gr2")

links = as.data.frame(matrix(c(0, 8, 21,
                               0, 9, 18,
                               0, 11, 8,
                               1, 8, 9,
                               2, 6, 5,
                               2, 7, 5,
                               2, 10, 11,
                               3, 8, 89, # cccc to cccc
                               3, 9, 13, # cccc to duuu
                               3, 12, 31, # cccc to not.detected
                               4, 9, 12, # dddu to duuu
                               5, 8, 29),
                             byrow = TRUE,
                             ncol = 3))

names(links) = c("source", "target", "value")

links$linkgroups <- c("gr3","gr4","gr5","gr6","gr7","gr8","gr9","gr10","gr11","gr12","gr13","gr14")

node_color <- 'd3.scaleOrdinal()
              .domain(["gr1", "gr2", "gr3", "gr4",
                       "gr5", "gr6", "gr7", "gr8",
                       "gr5", "gr6", "gr7", "gr8"])
              .range(["#C65911", "#FFE699", "#009ACD", "#00B2EE",
                      "#00BFFF", "#FFA500", "#CDAD00", "#EEC900",
                      "#FFD700", "#6E8B3D", "#A2CD5A", "#CAFF70",
                      "#FF8C69", "#CDC5BF"])'

p = sankeyNetwork(Links = links,
                  Nodes = nodes,
                  Source = "source",
                  Target = "target",
                  Value = "value",
                  NodeID = "name",
                  fontSize = 20,
                  nodeWidth = 100,
                  colourScale = node_color,
                  LinkGroup = "linkgroups",
                  NodeGroup = "nodegroups",
                  iterations = 0)
p
